# Global population structure and genotyping framework for genomic surveillance of the major dysentery pathogen, *Shigella sonnei*

**DOI:** 10.1101/2020.10.29.360040

**Authors:** Jane Hawkey, Kalani Paranagama, Kate S. Baker, Rebecca J. Bengtsson, François-Xavier Weill, Nicholas R. Thomson, Stephen Baker, Louise Cerdeira, Zamin Iqbal, Martin Hunt, Danielle J. Ingle, Timothy J. Dallman, Claire Jenkins, Deborah A. Williamson, Kathryn E. Holt

## Abstract

*Shigella sonnei* is the most common agent of shigellosis in high-income countries, and causes a significant disease burden in low- and middle-income countries. Antimicrobial resistance is increasingly common in all settings. Whole genome sequencing (WGS) is increasingly utilised for *S. sonnei* outbreak investigation and surveillance, but comparison of data between studies and labs is challenging. Here, we present a genomic framework and genotyping scheme for *S. sonnei* to efficiently identify genotype and resistance determinants from WGS data. The scheme is implemented in the software package Mykrobe and tested on thousands of genomes. Applying this approach to analyse >4,000 *S. sonnei* isolates sequenced in public health labs in three countries identified several common genotypes associated with increased rates of ciprofloxacin resistance and azithromycin resistance, confirming intercontinental spread of highly-resistant *S. sonnei* clones and demonstrating the genomic framework can facilitate monitoring of the emergence and spread of resistant clones at local and global scales.

## Introduction

*Shigella* spp are Gram-negative bacterial pathogens that cause shigellosis (bacterial dysentery). *Shigella* are transmitted via the fecal-oral route and estimated to cause ~188 million infections annually, leading to ~160,000 deaths mainly in young children^1^. In low- and middle-income settings, most of the *Shigella* disease burden of shigellosis is in children under five years^2^, however in high-income countries *Shigella* is frequently detected in returned travelers or men who have sex with men (MSM)^3,4^. *Shigella sonnei* is the most frequently isolated agent of shigellosis in high-income countries and in those that are economically developing^1,5,6^. *S. sonnei* emerged recently (~350 years ago^7^), share a single serotype, and display limited genomic diversity (all belong to ST152 complex by multi-locus sequence typing (MLST)). These properties make it difficult to differentiate and track *S. sonnei* strains^7^, motivating adoption of whole-genome sequencing (WGS) for research and public health surveillance of this organism^8^. Core-genome MLST (cgMLST) is available via the *Escherichia coli* scheme^9^ but has not been widely adopted for *S. sonnei* surveillance, and most public health labs and research studies rely on the higher-resolution technique of single nucleotide variant (SNV)-based phylogenetics analysis.

The global population of *S. sonnei* is divided into five major lineages^7,9^. Several WGS studies have investigated regional *S. sonnei* epidemiology and population structure, including in Asia^10–13^, Australia^4^, the Middle East^14^, South America^9^, and the United Kingdom^15–17^; and defined additional sub-lineage-level phylogenetic groups of local epidemiological importance, associated with features such as ciprofloxacin-resistance^18^, transmission within Orthodox Jewish communities^14^, or transmission amongst MSM^4,16^. *S. sonnei* from Asia, Europe, Australia and North America have for >20 years been dominated by Lineage 3 strains that are resistant to early first-line antimicrobials (trimethoprim-sulfamethoxazole, tetracycline, and streptomycin) due to the presence of antimicrobial resistance (AMR) genes acquired horizontally via the small plasmid spA and a chromosomal Tn*7*-like transposon^7,19,20^. Resistance to chloramphenicol and/or ampicillin is also observed (e.g. via acquisition of the *Shigella* resistance locus (SRL)^9,21^). Reduced susceptibility to fluoroquinolones has emerged on multiple occasions and in multiple locations through acquisition of point mutations within the quinolone resistance determining region (QRDR) of *gyrA*^7,10^. Resistance to ciprofloxacin has emerged at least once via the accumulation of three QRDR mutations (2 in *gyrA* and one in *parC*) in a South Asian sublineage that has since been detected on multiple continents^11–13,22^. Resistance to the last few remaining drugs is increasing through the acquisition and maintenance of plasmids carrying *mph(A)* and *ermB* (azithromycin resistance) or extended-spectrum beta-lactamase (ESBL) genes (ceftriaxone resistance), often in combination with additional aminoglycoside resistance genes^10,13,15,17,23^.

In many countries, *S. sonnei* is a notifiable infection and subject to public health surveillance and outbreak investigations, which are increasingly conducted using WGS^8,24–27^. However, the lack of a defined global genomic framework and accompanying genotype nomenclature hampers both local reporting, outbreak detection, and patterns of spread within regions. For example, most *S. sonnei* WGS studies have reported which of the five major lineages their novel isolates belong to, but have had to download public reference genome data, construct whole genome alignments, and infer phylogenies to achieve this basic identification^22,26^. Studies of MSM *S. sonnei* in different settings have designated different names for the same lineages^4,15,17^, obscuring the fact that the same clones are spreading amongst MSM communities in different countries, and the only way to recognize this currently is through construction of whole-genome phylogenies incorporating data from multiple prior studies^28^.

WGS-based genotyping frameworks based on single nucleotide variants (SNVs) have been widely adopted for the bacterial pathogens *Mycobacterium tuberculosis*^29^ and *Salmonella enterica* serovar Typhi^30^, which display similarly low levels of genomic diversity to *S. sonnei*. These frameworks enable fast and accurate typing of clinical isolates from WGS data without the need for time-consuming comparative genomics or phylogenetics, facilitating straightforward identification of (and cross-jurisdictional communication about) epidemiologically important lineages from WGS data.

Here, we describe the global population structure for *S. sonnei* and (i) propose a hierarchical SNV-based genotyping scheme, which we define using 1,935 globally distributed genomes; (ii) implement the scheme within the free and open-source Mykrobe^31^ software alongside detection of genetic determinants that are highly predictive of AMR phenotypes in *S. sonnei*^27^; and (iii) validate this approach to genotyping using an additional 2,015 genomes that were sequenced in public health laboratories and deposited in the publicly available GenomeTrakr database. By applying this novel genotyping framework to *S. sonnei* WGS data generated in public health laboratories on three continents, we demonstrate the utility of the new scheme for identifying, tracking and reporting emerging AMR clones both within and between jurisdictions.

## Results

### Defining phylogenetically informative genotypes for *S. sonnei*

In order to define the global population structure and identify clades and marker SNVs, we collated 1,935 publicly available *S. sonnei* genomes from eight studies^4,7,9–12,14,15^ as our ‘discovery’ dataset (see **Supplementary Tables 1 & 2**), in order to define the global population structure and identify clades and marker SNVs. These genomes represent isolates from 48 countries, collected between 1943 and 2018 (**Figure 1c-d**, **Table 1**). The majority originate from Asia (32.4%), Europe (29.3%), Australia (18.8%), or Latin America and the Caribbean (17.4%) (**Table 1**). The data set is diverse in terms of acquired AMR genes (median 9 per genome, range 0-21), and includes 150 (7.8%) genomes known to be associated with MSM.

**Figure 1:**
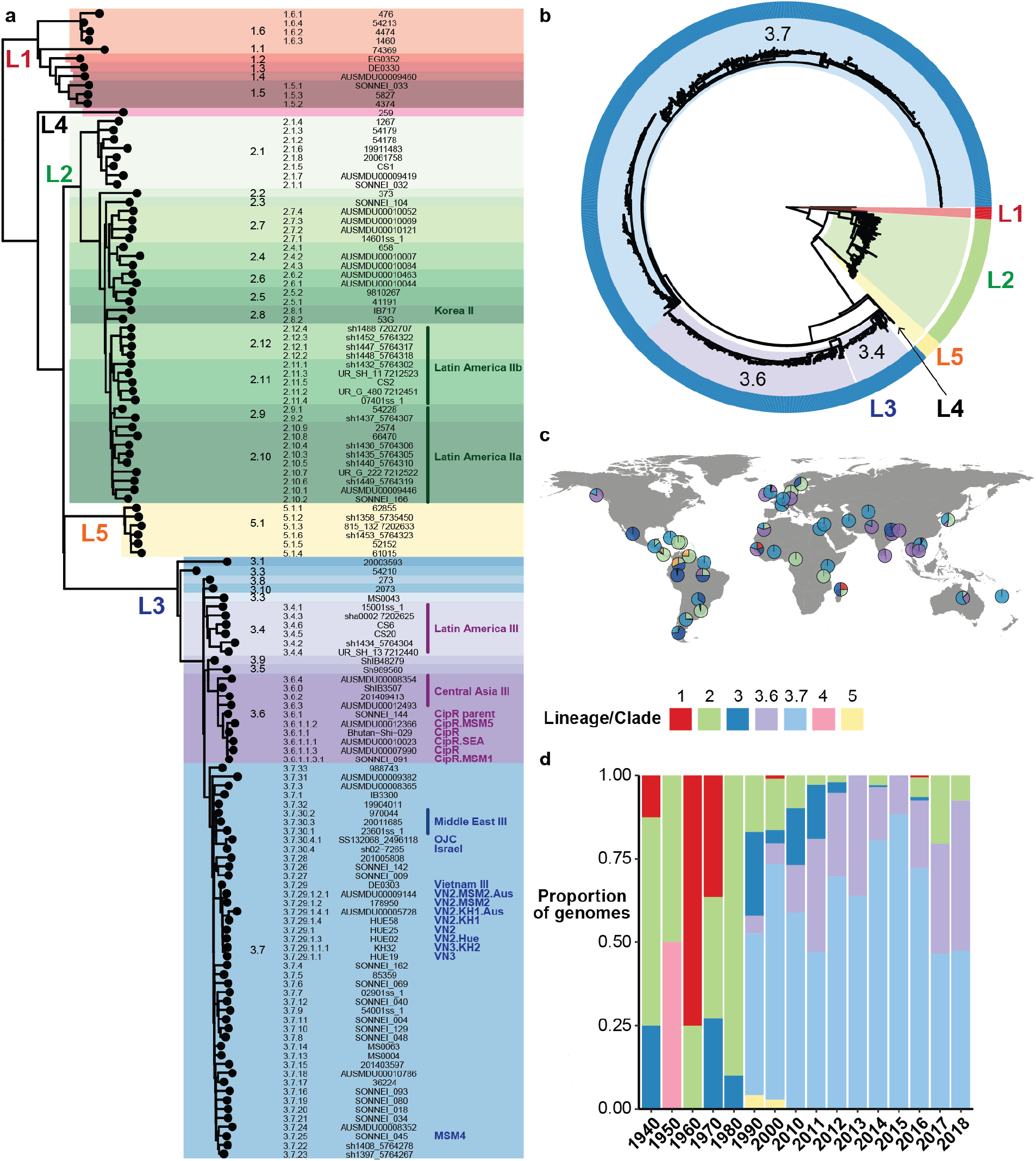
Population structure, temporal distribution and geographic distribution of the 1,935 *S. sonnei* genomes in the discovery set. **(a)** Maximum likelihood phylogeny (outgroup rooted using *E. coli*) of one representative per genotype. Lineages are labelled LX, where X is the lineage number. Highlighting and column 1 indicate clades, column 2 indicates genotype, column 3 shows strain names, column 4 shows human readable genotype names (for epidemiological groups noted in **Table 2**). **(b)** Maximum likelihood phylogeny (outgroup rooted using *E. coli*), **(c)** frequencies by geographic region, and **(d)** frequencies by decade/year; for all discovery set genomes and coloured by lineage and major clades (3.6, 3.7). Interactive version of linked phylogeny, map and timeline for this data set are available online in Microreact (https://microreact.org/project/fG2N7huk9oZNCaVHu8rukr).

**Table 1:** Geotemporal distribution of 1,935 *S. sonnei* genomes in the discovery dataset.

|  | Region | Country | Year(s) of isolation | No. of isolates |
| --- | --- | --- | --- | --- |
| Africa | **Total** |  | **1967-2006** | **19 (0.98%)** |
|  | Central Africa | Cameroon | 1973 | 1 |
|  | East Africa | Madagascar | 1998-2000 | 3 |
|  |  | Kenya, Tanzania | 2004 | 2 |
|  | Northern Africa | Egypt | 2005-2006 | 4 |
|  |  | Morocco | 2005-2006 | 4 |
|  | West Africa | Senegal | 1967-2006 | 4 |
|  |  | Burkina Faso | 2006 | 1 |
| Asia | **Total** |  | **1979-2014** | **627 (32.4%)** |
|  | Central Asia | Uzbekistan | 2005 | 1 |
|  | Eastern Asia | Korea | 1979-2003 | 20 |
|  | Southern Asia | Bhutan | 2011-2013 | 71 |
|  |  | India | 2013-2014 | 24 |
|  |  | Pakistan | 2002-2003 | 7 |
|  |  | Other (Nepal, Iran, Sri Lanka) | 2003-2006 | 3 |
|  | Southeast Asia | Cambodia | 2013-2014 | 4 |
|  |  | Thailand | 1994-2013 | 9 |
|  |  | Vietnam | 1995-2015 | 266 |
|  | Western Asia | Israel | 1992-2014 | 222 |
| Europe | **Total** |  | **1943-2016** | **567 (29.3%)** |
|  | Northern Europe | United Kingdom | 1990-2016 | 393 |
|  |  | Ireland | Unknown | 5 |
|  |  | Sweden | 1943-1947 | 6 |
|  |  | Denmark | 1945 | 1 |
|  | Western Europe | France | 1945-2014 | 158 |
|  |  | Belgium | 2008 | 3 |
|  |  | Germany | Unknown | 1 |
| Northern America | **Total** |  | **1994-2015** | **16 (0.8%)** |
|  | North America | USA | 2004-2015 | 13 |
|  |  | Other (unknown) | 1994-1995 | 3 |
| Latin America and the Caribbean | **Total** |  | **1997-2014** | **337 (17.4%)** |
|  | Caribbean | Cuba, Dominican Republic, Haiti | 2003-2006 | 3 |
|  | Central America | Costa Rica | 2002-2010 | 50 |
|  |  | Guatemala | 2011-2012 | 30 |
|  |  | Mexico | 1998 | 1 |
|  | South America | Argentina | 2002-2011 | 50 |
|  |  | Brazil | 1997-2002 | 7 |
|  |  | Chile | 2010-2011 | 27 |
|  |  | Colombia | 2008-2011 | 30 |
|  |  | French Guiana | 1998-2006 | 3 |
|  |  | Paraguay | 2008-2012 | 18 |
|  |  | Peru | 1999-2012 | 49 |
|  |  | Uruguay | 2000-2011 | 28 |
|  |  | Venezuela | 1997-2014 | 41 |
| Australia and Oceania | **Total** |  | **1997-2018** | **364 (18.8%)** |
|  | Australia | Australia | 2016-2018 | 363 |
|  | Oceania | New Caledonia | 1997 | 1 |
| Unknown | **Total** |  | **Unspecified** | **5 (0.5%)** |

The recombination-filtered core-genome maximum likelihood phylogeny inferred from these genomes (**Figure 1b**) was robust (median bootstrap support 100%) and exhibited the five previously-described deep branching lineages^7,9^. Lineage 3 was most common (86.9%), followed by Lineage 2 (10.7%), Lineage 5 (1.4%), Lineage 1 (0.9%) and Lineage 4 (n=1). The pairwise core-genome SNV distance distribution revealed peaks and troughs which we used to set thresholds to define clusters at different levels of resolution (**Figure 2a**). A threshold of 600 pairwise SNVs separated the five major lineages; troughs at 215 SNVs and 100 SNVs were used to define higher-resolution genetic clusters. (A similar structure was recovered using hierarchical Bayesian clustering of the SNV matrix using FastBAPS, but with less consistent levels of divergence between clusters; see **Supplementary Figure 1**). Mapping the pairwise SNV threshold-defined clusters onto the phylogeny confirmed that each cluster corresponded to a monophyletic group with 100% bootstrap support, which we designate as clades (n=29, using 215-SNV threshold) and subclades (n=96, using 100-SNV thresholds).

**Figure 2:**
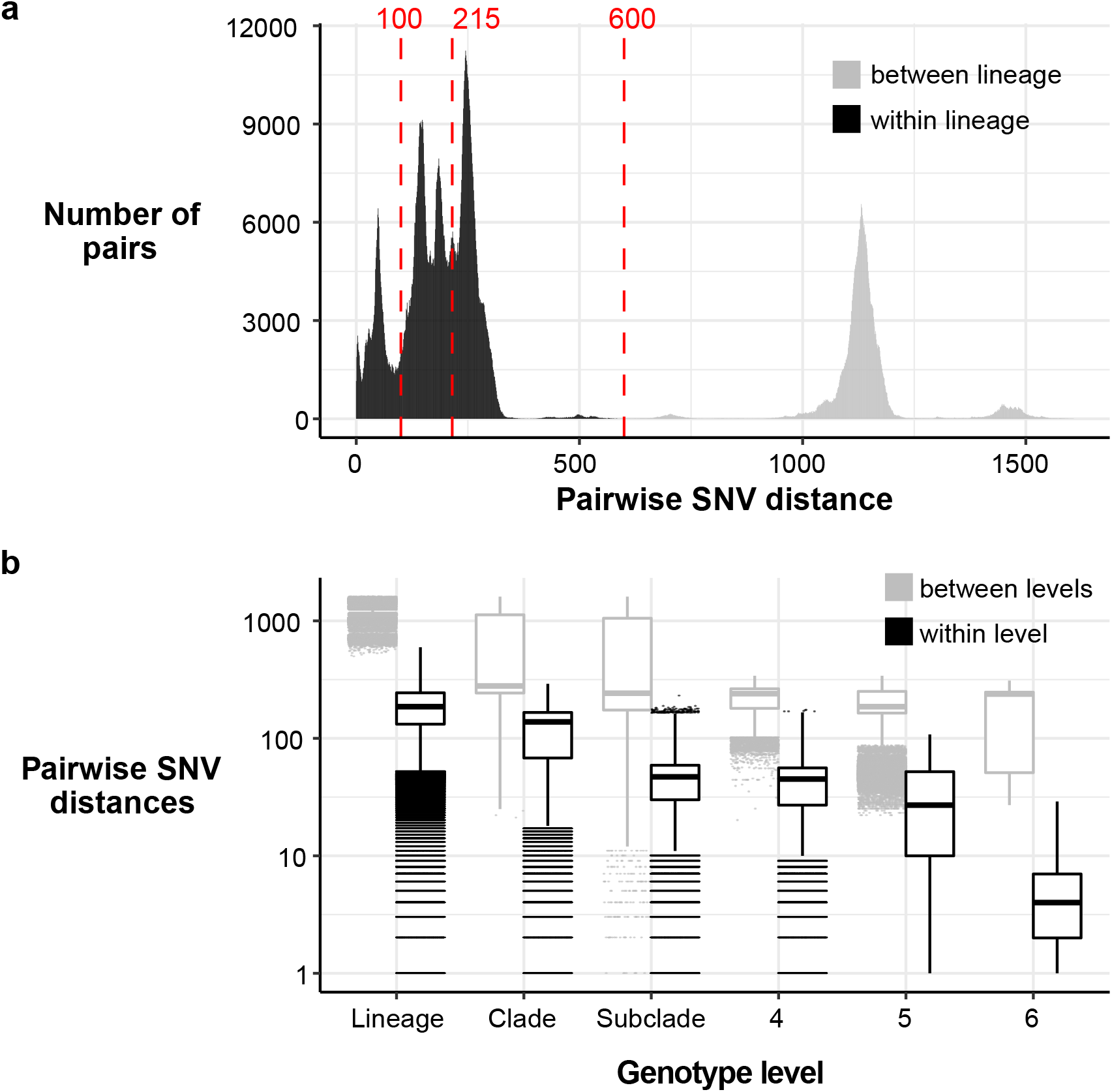
Distribution of SNV distances in discovery genomes. **(a)** Histogram of pairwise SNV distances between all discovery genomes, coloured by lineage comparison as per legend. Red lines mark SNV cut-offs used to define lineage, clade and subclade levels in genotyping scheme. **(b)** Boxplots of pairwise SNV distances (log scale) between discovery genomes at different levels of the defined genotyping scheme.

We used these cluster memberships to define hierarchical genotypes with nomenclature in the form [lineage].[clade].[subclade] (see **Figure 1a**). Similar to the *M. tuberculosis* and *S*. Typhi schemes, this hierarchical nomenclature facilitates easy recognition of relationships between genotypes; e.g. subclades 3.6.1, 3.6.2,…, 3.6.N are sister groups in the whole-genome phylogeny, nested within clade 3.6, which falls within Lineage 3 (see **Figure 1a**). The median pairwise distance between genomes of the same clade or subclade was 138 or 47 core-genome SNVs, respectively (**Figure 2b**).

Whilst the discovery set is not a systematic sampling across geographic regions, it can provide some preliminary insights into the global distribution of *S. sonnei* genotypes. Lineages were broadly distributed across continents (with the exception of Lineage 5 and the singleton Lineage 4, see **Figure 1c**), however the majority of clades (n=24, 83%) were represented by isolates from just one (n=16, 57%) or two (n=7, 23%) continents. At the other extreme, clades 3.6 and 3.7 were widely distributed, with representatives on all six continents (**Figure 1b**). Clades 3.6 and 3.7 were the most common overall (19% and 62%, respectively) and accounted for the majority of *S. sonnei* from all continents except Latin America, where clades 2.10, 2.11, 2.12, 3.4, 3.6 and 5.1 were common (8-28% each). Subclades showed even greater geographic specificity, with 74% (n=71) represented by a single continent only and 72% (n=69) represented by a single region only. Fifty-nine subclades (61%) were dominated by genomes from a single country.

As a key goal of the *S. sonnei* genotyping scheme is to facilitate identification and communication about subtypes of public health interest, we reviewed the position of genetic clusters that have been described in the literature as being of epidemiological importance (**Table 2**). Groups previously identified as being associated with specific geographical regions mapped mainly to clades or subclades defined in the genotyping scheme (**Table 2**). Most groups previously defined on the basis of AMR or transmission patterns comprised more recently-emerged clusters, forming monophyletic groups within our subclade-level genotypes. Hence we created additional higher-resolution genotypes nested within subclades to demarcate these groups (e.g. 3.6.1.1, 3.6.1.1.1; see **Table 2**), and anticipate adding more genotypes as new resistant groups emerge in future. For example, the ciprofloxacin resistant triple-mutant sublineage^11–13^ comprised a monophyletic group within subclade 3.6.1 that we define as genotype 3.6.1.1; distinct subgroups within this have also been described, associated with South East Asia (genotype 3.6.1.1.1), and MSM communities in Australia (3.6.1.1.2) or the UK (3.6.1.1.3.1) (see **Table 2**).

**Table 2:** Details of epidemiological clusters defined within the *S. sonnei* population

| New genotyping framework |  | Previously defined as |  | Description |
| --- | --- | --- | --- | --- |
| Genotype | Name | Name | Study |  |
| Lineage II |  |  |  |  |
| 2.8 | Korea II | Korea II | Holt 2012 | Associated with Korea |
| 2.9, 2.10, 2.11 | Latin America II | South America II | Holt 2012 | Associated with Latin America |
|  |  | LA sublineage IIa & IIb | Baker 2017 |  |
| Lineage III |  |  |  |  |
| 3.4 | Latin America III | South America III | Holt 2012 | Associated with Latin America |
|  |  | LA sublineage IIIa & IIIb | Baker 2017 |  |
| 3.6 | Central Asia III | Central Asia IIIa | Holt 2012 | Associated with Central Asia |
| 3.6.1 | CipR parent | - | This study | Subclade from which ciprofloxacin-resistant sublineage emerged |
| 3.6.1.1 | CipR | Ciprofloxacin-resistant | The 2015 | Ciprofloxacin-resistant triple mutation sublineage |
|  |  | Pop2 | The 2019 |  |
| 3.6.1.1.1 | CipR.SEA | - | This study | Ciprofloxacin-resistant isolates associated with South East Asia |
| 3.6.1.1.3.1 | CipR.MSM1 | MSM clade 1 | Baker 2018 | MSM-linked ciprofloxacin resistant isolates |
| 3.6.1.1.2 | CipR.MSM5 | BAPS1 | Ingle 2019 | MSM-linked ciprofloxacin resistant isolates |
|  |  | MSM Clade 5 | Bardsley 2020 |  |
| 3.7.25 | MSM4 | MSM Clade 4 | Baker 2018 | MSM-linked |
| 3.7.29 | Vietnam III | VN clone | Holt 2012 | Associated with South East Asia |
| 3.7.29.1 | VN2 | VN clone, sweep 2 | Holt 2013 | Clonal group originating from genetic sweep 2 |
| 3.7.29.1.1 | VN3 | VN clone, sweep 3 | Holt 2013 | Clonal group originating from genetic sweep 3 |
| 3.7.29.1.1.1 | VN3.KH2 | KH2 | Holt 2013 | Kanh Hoa subclone 2, emerging from sweep 3 |
| 3.7.29.1.1.2 | VN4 | VN clone, sweep 4 | Holt 2013 | Clonal group originating from genetic sweep 4 |
| 3.7.29.1.2 | VN2.MSM2 | MSM Clade 2 | Bardsley 2020 | MSM-linked strains, emerging from sweep 2 |
| 3.7.29.1.2.1 | VN2.MSM2.Aus | - | This study | Australian MSM-linked, emerging from sweep 2 |
|  |  | (part of BAPS3) | Ingle 2019 |  |
| 3.7.29.1.3 | VN2.Hue | Hue2 | Holt 2013 | Hue subclone 2, emerging from sweep 3 |
| 3.7.29.1.4 | VN2.KH1 | KH1 | Holt 2013 | Kanh Hoa subclone 1, emerging from sweep 2 |
| 3.7.29.1.4.1 | VN2.KH1.Aus | - | This study | Australian isolates, emerging from KH1 |
|  |  | (part of BAPS3) | Ingle 2019 |  |
| 3.7.30 | Middle East III | Middle East III | Holt 2012 | Associated with Middle East |
| 3.7.30.4 | Israel III | Israel III | This study | Associated with Israel |
| 3.7.30.4.1 | OJC | OJC-associated | Baker 2016 | Associated with the Orthodox Jewish communities in Israel, UK, USA and Europe |

To facilitate communication about genotypes of epidemiological interest, we also assigned them human readable aliases (e.g. 3.6.1.1 = CipR, 3.6.1.1.1 = CipR.SEA, 3.6.1.1.2 = CipR.MSM5, see **Table 2**). As far as possible these aliases map to names given in previous publications, e.g. the MSM clade numbers designated in^15,16^. Most of the epidemiological groups of interest belong to Lineage 3, and detailed phylogenies for these groups are provided in **Supplementary Figure 2**. In addition to being monophyletic on the tree, these higher-level genotypes were supported by FastBAPS analysis (**Supplementary Figure 1**). Pairwise distances within and between genotypes of all levels are shown in **Figure 2b**.

### Development and validation of SNV-based scheme for assigning genotypes

We identified marker SNVs unique to each genotype (147 SNVs in total, see **Supplementary Table 3**) and implemented code to assign new genomes to genotypes based on presence of these markers (see **Methods**, available in Mykrobe v0.9.0, https://github.com/Mykrobe-tools/mykrobe). To validate this approach, we downloaded and genotyped 2,015 additional *S. sonnei* genomes from GenomeTrakr (referred to as validation set, see **Supplementary Table 2**). These genomes originate from public health laboratories in three countries (n=609 USA, n=1,325 UK, n=11 Israel, n=70 country unknown), with isolation dates between 2015 and 2019.

We identified 17 different genotypes, all belonging to clades 3.6 or 3.7 (**Supplementary Table 4**). The vast majority (70%, n=1,403) belonged to clade 3.6. Genotype 3.6.1.1.2 (CipR.MSM5) was the most prevalent, assigned to 26.6% of the genomes, followed by 3.6.1.1 (CipR, 19.6%) (**Supplementary Table 4**). The UK GenomeTrakr genomes yielded the greatest number of genotypes (n=16), followed by the USA (n=13); likely due to a high number of travel-associated cases. All GenomeTrakr genomes deposited from Israel were identified as 3.7.30.4 (Israel III, 9%) or 3.7.30.4.1 (OJC, 91%); these genotypes were also detected amongst UK and USA genomes.

To verify the genotyping scheme accurately captured the population structure present in the GenomeTrakr isolates, we constructed a core-genome phylogeny including both the validation set and Lineage 3 discovery set (total n=3,696 genomes, see **Methods**) and mapped the genotype assignments to this tree (see **Supplementary Figure 3** and Microreact https://microreact.org/project/g8BvA2JCXWaZNDyPyjsWXF). All groups of isolates sharing a genotype assignment based on marker SNVs constituted monophyletic clades within the core-genome phylogeny, consistent with the expected behavior of the scheme. This was true for all levels in the hierarchical scheme, including clades, subclades, and higher-resolution epidemiological groups.

### Distribution of antimicrobial resistance determinants amongst *S. sonnei* genotypes

We used the genotyping scheme to facilitate exploration of the distribution of AMR determinants in the global *S. sonnei* population, by assessing the frequency of AMR genes and QRDR SNVs across genotypes (**Figure 3**). For this analysis we included n=6,715 genomes: n=1,935 discovery set, n=2,015 validation set and a further n=2,765 public genomes (accessions listed in **Supplementary Table 2**). Most AMR determinants were associated with specific genotypes, present amongst either all or no members of each genotype (**Supplementary Figure 4**).

**Figure 3:**
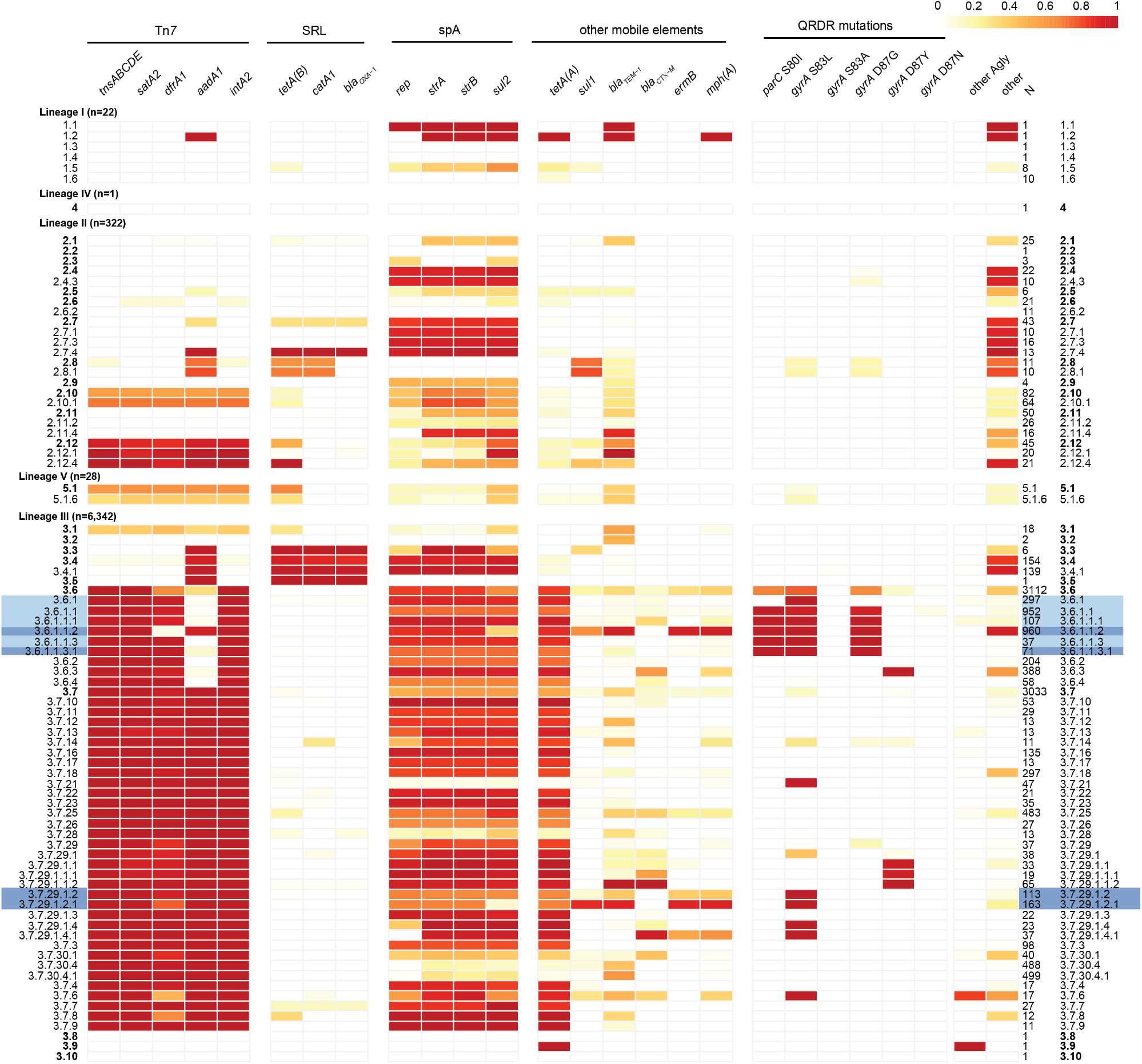
Frequencies of AMR genetic determinants within individual *S. sonnei* genotypes, calculated across 6,715 genomes. Cells indicate absence (white) or presence (coloured by proportion as per legend) of each AMR determinant (columns) within each clade or higher-resolution genotype (rows). All clades are included as rows (bold labels); subclades higher-resolution genotypes represented by ≥10 genomes are also included as distinct rows; number of genomes in each row are noted in column ‘N’. Light blue shading indicates fluoroquinolone resistant genotypes; dark blue shading indicates MSM-associated genotypes. Columns are grouped by typical location of the AMR determinant (labelled horizontal bars at the top): transposon Tn*7*, represented by marker genes *tnsABCDE* and class II integron In*2* integrase gene *intA2; Shigella* resistance locus (SRL); spA plasmid, represented by marker gene *rep;* other mobile elements; mutations in quinolone resistance determining region (QRDR). Column ‘other Agly’ indicates proportion of genomes carrying at ≥1 additional aminoglycoside resistance gene beyond those with their own columns; column ‘other’ indicates proportion of genomes carrying ≥1 other AMR gene that is not otherwise listed (full AMR gene content per strain is available in **Supplementary Table 1**).

Genes conferring resistance to first-line drugs were found in all lineages. Those associated with the spA plasmid (*sul2, tetA(A)*) were found in all lineages but were most widely distributed across clades of Lineage 3 (found in all clades) followed by Lineage 2 (81% of clades) (**Figure 3, Supplementary Figure 4**). Tn*7* transposon genes (*tnsABCDE*), the class II integron integrase (*intA2*) and AMR genes in the integron cassette (*satA2, dfrA1, aadA1*) were absent from Lineage 1 but found in Lineages 2 and 3 (40-41% of clades) and the single Lineage 5 clade (**Figure 3**). This combination of markers for the chromosomally integrated MDR transposon Tn*7* was most common in clades 3.6 (99%), 3.7 (99%) and 2.12 (88%), where it was typically accompanied by spA genes *(sul2, tetA(A))* resulting in resistance to co-trimoxazole (**Figure 3**). First-line AMR genes *aadA1, tet(B), catA1* and *bla*_OXA-1_, which are known to mobilise together on the SRL, co-occurred in clades 3.3, 3.4 and 3.5, consistent with prior reports of SRL in clade Latin America IIIa (3.4)^9^. Acquired genes *bla*_TEM-1_ and *sul1* were also found at low frequencies across diverse genotypes (**Supplementary Figure 4**), suggesting occasional acquisition via mobile elements.

QRDR SNVs were detected in 50.4% of all genomes, distributed across six clades and 12 subclades (**Figure 3**), consistent with frequent emergence of these mutations under selection from drug exposure. Most common was GyrA-S83L (42% of genomes, six clades, 12 subclades) followed by GyrA-D87G (30.8% of genomes, five clades, ten subclades). Single mutants were most common (18.7% of genomes, five clades, 6 subclades) but double mutants were also observed in 3.6.1 (n=61/6,715 genomes had GyrA-S83L + GyrA-D87G). QRDR triple mutants, associated with ciprofloxacin resistance, were detected only in the CipR sublineage (genotype 3.6.1.1) which harbours GyrA-S83L + GyrA-D87G + ParC-S80I. The emergence and evolutionary dynamics of this CipR sublineage from within the Central Asia IIIa clade (genotype 3.6) were recently described in a detailed phylodynamic study of fluoroquinolone resistant isolates from diverse sources by The *et al*^13^. That study divided the Central Asia III clade into two populations: Pop1, with either GyrA-D87Y or GyrA-S83L arising on two independent occasions in South Asia in the mid-1990s; and Pop2, which arose from Pop1 genomes carrying GyrA-S83L in South Asia in the early 2000s, and then acquired GyrA-D87G and ParC-S80I to become fluoroquinolone resistant before spreading geographically^13^. Applying our new genotyping scheme to the genomes from The *et al*^13^ (**Supplementary Figure 5**), we confirm that Pop1 maps to clade 3.6 (n=18) and its subclades 3.6.1, 3.6.2, 3.6.3, 3.6.4; and Pop2 maps to sublineage 3.6.1.1 (CipR, n=239) including its subgroups 3.6.1.1.1 (CipR.SEA, n=30), 3.6.1.1.2 (CipR.MSM5, n=3), 3.6.1.1.3 (n=16) and 3.6.1.1.3.1 (CipR.MSM1, n=19).

Determinants of resistance to azithromycin and extended-spectrum cephalosporins were rare and concentrated mainly in clades 3.6 and 3.7. The plasmid-borne azithromycin resistance genes *mph(A)* and *ermB* were detected at high frequency in genotypes 3.6.1.1.2 (CipR.MSM5, n=915, 95%) and 3.7.29.1.2.1 (VN2.MSM2.Aus, n=147, 90%); *mph(A)* was present alone in the single 1.2.1 genome (Vietnam, 2007), and alone or with *ermB* at lower frequency (<65%) amongst other Lineage 3 genotypes (**Figure 3**). Notably, n=947 3.6.1.1 (CipR) genomes carried *mph(A)* in addition to Tn*7* and spA genes, rendering them resistant to azithromycin, ciprofloxacin and first-line drugs (n=58, 3.6.1; n=16, 3.6.1.1; n=868, 3.6.1.1.2; n=2, 3.6.1.1.3; n=3, 3.6.1.1.3.1), leaving extended-spectrum cephalosporins as the last remaining oral drug. ESBL genes were detected only sporadically, at low frequencies in clades 3.6 (13%) and 3.7 (12%), across 25 distinct genotypes (frequency range, 0.6-100%, median 17.2%) (see **Figure 3**). Carbapenemase genes were extremely rare, present in only two genomes (*bla*_OXA-66_ and *bla*_OXA-181_). Concerningly, we detected 40 genomes with resistance determinants for azithromycin, third-generation cephalosporins and fluoroquinolones, all within CipR genotype 3.6.1.1 (n=16, 3.6.1.1; n=11, 3.6.1.1.1; n=15, 3.6.1.1.2). These genomes were isolated between 2014-2019, and were found in genomes from England (n=21), Australia (n=13), the USA (n=3), Vietnam (n=3) and the Netherlands (n=2) (further details below).

### Application to public health surveillance data from Australia, England and USA

To demonstrate how the *S. sonnei* genotyping framework can facilitate the rapid tracking and reporting of emerging AMR trends across jurisdictions, we applied it to genomic surveillance data from Victoria, Australia (n=644), England (n=2,867) and USA (n=711) generated over a 4-year period (2016 to 2019) (data in **Supplementary Table 2**). The data represents all cultured isolates submitted to the Microbiological Diagnostic Unit Public Health Laboratory in Australia (42% of all *S. sonnei* notifications in Victoria)^4^ and all those sent to the Public Health England Gastrointestinal Bacteria Reference Unit^27^ (provided direct from the reference laboratories for the present study); and ~5% of those notified in the USA (sourced from the public GenomeTrakr database^32^). The total time taken to generate genotyping reports (including QRDR mutations) for all n=6,715 isolates was ~40 sec – 1 min per isolate using Mykrobe, with raw Illumina sequence files (fastq format) as input.

**Figure 4a** shows the annual frequency of fluoroquinolone resistance (defined as presence of 3 QRDR mutations) in each country, and the distribution of genotypes amongst the resistant isolates. Increasing fluoroquinolone resistance rates are evident amongst the *S. sonnei* samples from each country, beginning at ≤25% in 2016 and reaching 85% in Australia, 39% in England and 44% in USA in 2019 (black lines, **Figure 4a**). All fluroquinolone resistant genomes belonged to genotypes within the CipR sublineage (3.6.1.1), with the subgroup 3.6.1.1.2 (CipR.MSM5) accounting for a steadily increasing proportion of resistant isolates in each country, from ≤30% in 2016 to 78% in Australia, 59% in England and 85% in USA in 2019 (pink bars, **Figure 4a**). These results are consistent with local epidemiological outbreaks of *S. sonnei* in MSM communities in England^15,16^ and Australia^4,33^. Notably however, the common nomenclature makes it easy to identify several epidemiologically important patterns: (i) all resistant isolates in all three countries derive from the previously-described CipR sublineage 3.6.1.1 that emerged from South Asia in the early 2000s; (ii) the reported spread of resistant *S. sonnei* in MSM communities in Australia and England involves the same strain (this was not clear from previous reports, as the strain was named ‘MSM clade 5’ in the English studies and formed a subgroup within the ‘BAPS3 cluster’ in the Australian studies); (iii) this strain represents a clonal subgroup of the CipR sublineage (genotype 3.6.1.1.2, CipR.MSM5) that has disseminated intercontinentally over the last few years and become responsible for the majority of fluoroquinolone resistant *S. sonnei* infections in all three countries. (Note an additional two English isolates carried *gyrA*-S83L and *parC*-S80I plus the *qnrS* gene, which likely combine to confer fluoroquinolone resistance^34^).

**Figure 4:**
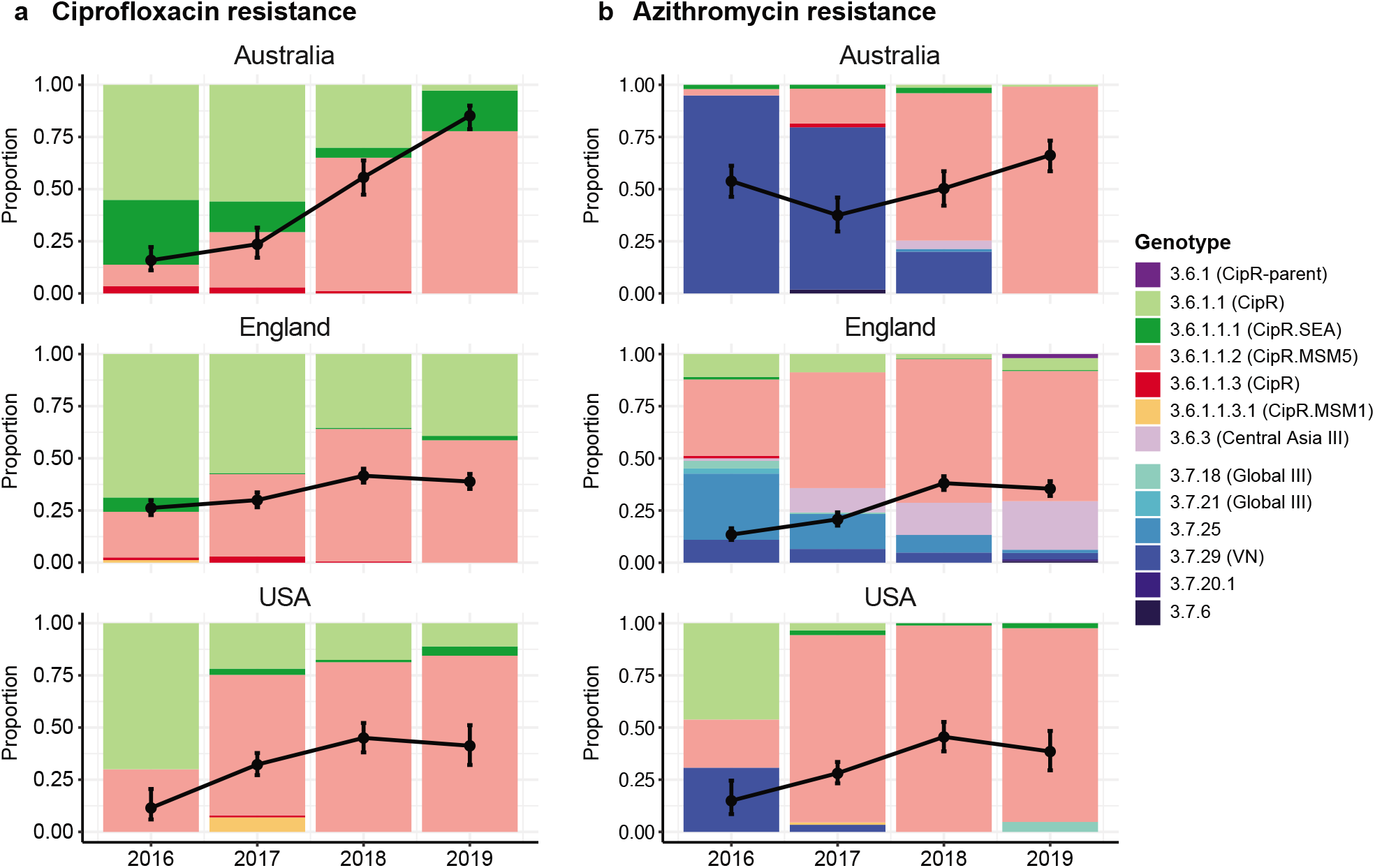
Prevalence and genotype breakdown of ciprofloxacin and azithromycin resistant *S. sonnei* in three surveillance regions (Australia, England and USA) from 2016 – 2019. In each plot, black lines indicate the proportion of genomes that are predicted resistant to **(a)** ciprofloxacin (defined as presence of ≥3 QRDR mutations) or **(b)** azithromycin (defined as carrying *mph(A))*; error bars indicate 95% confidence intervals for the proportion resistant. Stacked bars indicate the relative abundance of each genotype among resistant isolates.

**Figure 4b** shows the annual frequency of azithromycin resistance (predicted by presence of *mph(A))* in each country, and the genotypes responsible. Resistance rates were high (>50%) in Australia across the whole period, reflecting documented outbreaks in the MSM community^4^. In England and the USA, rates increased between 2016 and 2019, from 13% to 38% in England and 15% to 45% in the USA (black lines, **Figure 4b**). The genotype distributions amongst *mph(A)+* genomes differed markedly between countries in 2016, dominated by 3.7.29 (VNclone, 95%) in Australia, 3.7.25 (MSM4, 32%) and 3.6.1.1.2 (CipR.MSM5, 36%) in England, and 3.6.1.1 (CipR, 46%), 3.7.29 (VNclone, 31%) and 3.6.1.1.2 (CipR.MSM5, 23%) in the USA (see barplot, **Figure 4b**). Notably though, the contribution of 3.6.1.1.2 (CipR.MSM5) increased dramatically in each country, and in 2019 accounted for 99% of *mph(A)+* isolates in Australia, 62% in England and 93% in USA (bars, **Figure 4b**). Thus in 2019, the majority of predicted azithromycin resistant isolates in all three countries were also predicted to be resistant to ciprofloxacin and first line drugs. Concerningly, the proportion of total *S. sonnei* genomes with combined resistance determinants for azithromycin, ciprofloxacin and first-line drugs was high (36-66%) in all three countries in 2019 (**Figure 5**).

**Figure 5:**
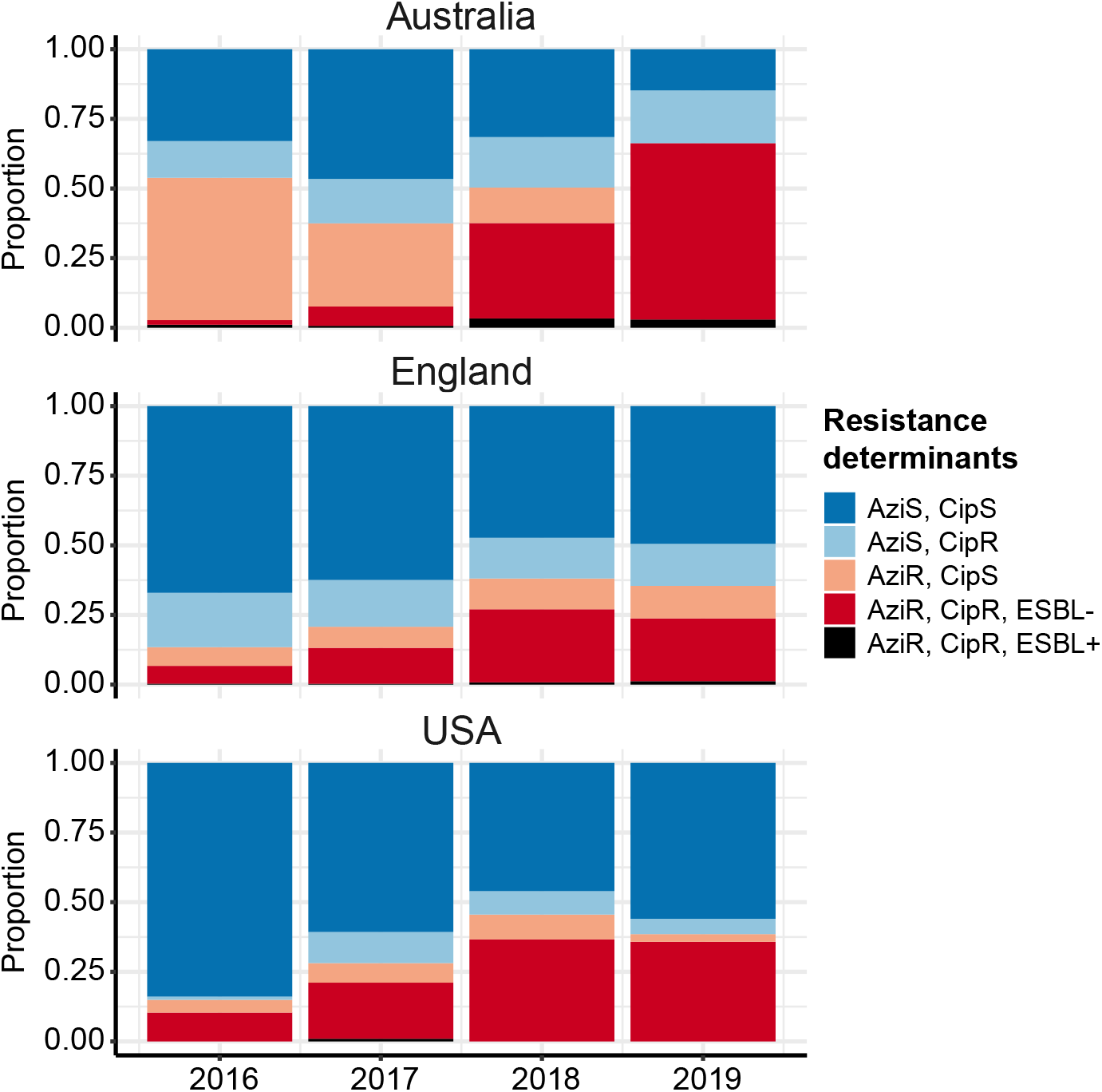
Prevalence of combined resistance to ciprofloxacin and azithromycin amongst genomes from each surveillance region. Stacked bar colours indicate the relative abundance of different combinations of resistances, predicted from genomes: CipR, ciprofloxacin resistant (defined as presence of ≥3 QRDR mutations); AziR, azithromycin resistant (defined as carrying *mph(A)*); ESBL+, presence of extended-spectrum beta-lactamase (ESBL) gene associated with resistance to third generation cephalosporins (*bla*_CTX-M-14_, *bla*_CTX-M-15_, *bla*_CTX-M-27_, *bla*_CTX-M-55_, *bla*_CTX-M-134_).

Whilst ESBL genes were rare across the *S. sonnei* surveillance data (7.8% in Australia, 17.5% in England, 1% in USA, see **Supplementary Table 2**), concerningly 201 genomes carried ESBL genes in addition to *mph(A)*, and 36 of these were QRDR triple mutants belonging to 3.6.1.1 (CipR). The latter were concentrated in genotypes 3.6.1.1, 3.6.1.1.1 and 3.6.1.1.2 and comprised 13 Australian genomes (2% of total, 2016-2019, all three genotypes), 20 English genomes (0.7% of total, in 2016 and 2018-2019, all three genotypes) and 3 USA genomes (0.4% of total, 2017, only 3.6.1.1 and 3.6.1.1.1) (**Figure 5**). Consistent with the overall picture of ESBL genes in *S. sonnei* described above (**Figure 3**), there was no evidence of strong linkage between specific ESBL genes and particular genotype backgrounds in the public health surveillance data for this period (**Supplementary Table 2**). This was true even amongst CipR/AziR/ESBL+ genomes, which included eight unique combinations of genotype and *bla*_CTX-M_ allele (3-6 per country, **Supplementary Figure 6**), consistent with multiple independent acquisitions of different AMR plasmids in different settings contributing to the march towards pan-resistance to oral drugs.

## Discussion

Here, we provide a global framework for *S. sonnei*, and identify marker SNVs that can be used to easily position newly sequenced isolates into this framework, without the need for time-consuming comparative genomics or phylogenetic analysis. We demonstrate that the population structure of *S. sonnei* can be represented by a robust maximum likelihood phylogeny and define within it 137 subtrees on the basis of pairwise divergence and epidemiological coherence, which we designate as hierarchically nested genotypes. Further, we provide a software package implemented within the Mykrobe^31^ code base, which can identify both the *S. sonnei* genotype and AMR determinants direct from short-read sequence files in a few seconds.

The genotyping scheme was constructed with a view towards stability, prioritising as markers SNVs that are found in highly conserved core genes under purifying selection in the population. We also endeavoured to make the scheme backwards compatible by identifying and designating unique genotypes for *S. sonnei* genetic clusters defined in previous studies on the basis of epidemiological features (see **Table 2**). We also aimed to ensure genotypes are interpretable, with stable numerical identifiers that convey relationships between genotypes, and human readable aliases that convey relevant epidemiological information where appropriate (see **Table 2**). The genotyping scheme can be readily expanded in future, by adding new genotypes and corresponding SNV markers as new clusters emerge. We envisage managing this via an international working group consisting of epidemiological, public health and genomics experts, similar to the approach proposed for a recent *Neisseria gonorrhoeae* typing scheme^35^. Importantly, our approach provides genotype definitions and nomenclature that are stable and transparent, not dependent on comparative analysis or on any specific sequencing assay or software package (although we provide an implementation in the Mykrobe v0.9.0 software for convenience, which has been tested on short and long reads (ONT) as input). This is designed to facilitate easy communication between laboratories and jurisdictions, and straightforward comparison of pathogen populations over time, allowing for rapid identification of clonal dissemination and AMR trends.

To demonstrate the utility of the genotyping scheme for public health applications, we applied it to summarise large genomic surveillance datasets from three jurisdictions in Australia, England and the USA. The results provide a straightforward view of temporal trends in the populations of *S. sonnei* causing disease in each jurisdiction. They also clearly identify common ciprofloxacin and azithromycin resistant clones that have spread globally and are now present in all three jurisdictions, which was previously not obvious from individual studies, as there was no common nomenclature.

The genotyping approach introduced here could greatly simplify the bioinformatics procedures required for routine genomic surveillance of *S. sonnei* in reference laboratories, the first step of which usually involves comparison of newly sequenced genomes to those from prior cases in the same jurisdiction (to monitor local trends) and/or other jurisdictions (to monitor introduction of new strains and patterns of regional spread). Notably, the public health labs in Australia, England and USA from which we sourced genomic surveillance data (**Figure 4**) all utilise genome-wide SNV-based phylogenetics for *S. sonnei* analysis^16,33^. An alternative approach to SNV-based analyses is cgMLST^36^, however this also requires comparison to a large database of other sequences; it also typically requires genome assembly, and the arbitrary nature of the assigned ST numbers makes it impossible to ascertain relationships between genotypes without database look-up or comparative analysis. Perhaps for this reason, we could identify only one report of *S. sonnei* analysis that utilised cgMLST^37^, and this study also relied on SNV-based phylogenetics to place their local isolates in the context of global populations (see Example 2 in **Supplementary Text, Supplementary Table 5**).

Reliance on whole genome comparisons and phylogenetic inference is considerably slower than genotyping and requires expertise and background knowledge both to conduct the analysis and to interpret the results. In contrast, using the genotyping framework, raw sequence data can be turned into simple informative identifiers for each strain without reference to other genomes or databases (and without needing to assemble the genome). The resulting genotyping information can be easily interpreted, compared and stored in (non-sequence) databases for future reference, facilitating epidemiological investigations without need for direct comparisons with any other genome sequences. Detailed phylogenetic analysis can then be focused on subsets of isolates that share the same or similar genotypes, if needed to address specific questions (e.g. relating to emerging local outbreaks or transmission networks).

For example, scientists investigating azithromycin resistant *S. sonnei* isolated from MSM in Switzerland recently reported identifying the strains as belonging to the same clones spreading through MSM communities in England. To achieve this identification, they had to download English genome data, compare their newly isolated genomes to these using read mapping, and construct and interpret whole genome phylogenies^28^. A similar study using WGS to investigate a ciprofloxacin resistant outbreak in California used the same informatics approach to conclude that the local strain belonged to the previously described ciprofloxacin resistant lineage originating in South Asia^22^. Another recent study from Switzerland used WGS to investigate an increase in ESBL *S. sonnei* using a combination of cgMLST and phylogenetics^37^. Using our new genotyping approach, all of these identifications could be made within minutes of obtaining sequence data, with no need for external comparative data, background knowledge of *S. sonnei* genetics, or complex computational infrastructure and expertise (see **Supplementary Text**, **Supplementary Figure 7**, **Supplementary Tables 5 & 6**). The rapid identification of genomically related isolates could be used to facilitate timely public health responses to shigellosis outbreaks.

In conclusion, while genomics is increasingly becoming a standard tool for surveillance of *S. sonnei* and other pathogens in public health reference laboratories, this poses computational and epidemiological challenges in terms of analysis, interpretation and communication of genome-derived data across discipline and jurisdictional boundaries. The genotyping framework and universal nomenclature for *S. sonnei* established here provides a solution for many of these issues. Importantly, it will facilitate monitoring of the emergence and spread of AMR *S. sonnei* clones, at local and global levels, which will become increasingly important as public health agencies face the emerging threat of pan-resistant *S. sonnei*.

## Methods

### Single nucleotide variants (SNVs) and phylogenetic analysis – discovery dataset

The discovery dataset consisted of 1,935 high quality previously published *S. sonnei* genomes, sequenced using Illumina platforms. Source information for all genomes (year of collection, geographic origin, etc) was extracted from their respective publications^4,7,9–12,14,15^ (**Supplementary Tables 1 & 2**). Genomes were mapped to the *S. sonnei* reference genome 53G (accession NC_016822) using RedDog (v1beta11; https://github.com/katholt/RedDog). Briefly, RedDog maps reads with Bowtie2 v2.2.9^38^ with the sensitive local parameter, then uses SAMtools v1.1^39^ to retain high quality SNV calls (phred score ≥20, read depth >5, removes heterozygous calls). SNVs detected in repetitive regions where SNV calls are dubious (e.g. insertion sequences, phage) were removed (see Figshare, https://doi.org/10.26180/5f1a443b19b2f). SNVs associated with recombination were identified and excluded using Gubbins v2.3.2^40^. The final alignment consisted of 23,673 SNVs across 1,935 genomes. Ancestral alleles at these SNV sites were extracted from five *E. coli* genomes (accessions CP019005, CP031916, CP019961, CP034399 and CP019259) using the mapping procedure described above, and were included in the alignment for the purpose of outgroup rooting the tree. A maximum likelihood (ML) phylogeny was inferred using IQ-TREE v2^41^ using a GTR substitution model (**Figure 1b**). An interactive form of the tree is available in Microreact, at https://microreact.org/project/fG2N7huk9oZNCaVHu8rukr.

### Defining clades and subclades of the genotyping scheme

The discovery set was analysed to define appropriate SNV thresholds for assigning genomes to genotypes based on pairwise genetic distances. Pairwise SNV distances were calculated for all pairs of, and thresholds were selected by examining the distribution of pairwise SNV distances (**Figure 2a**), to define clusters at three levels: lineage (600 SNVs), clade (215 SNVs) and subclade (100 SNVs). To cluster discovery set genomes at these thresholds, we applied hierarchical clustering (using *hclust* function in R, with complete linkage) to the pairwise SNV distance matrix. The resulting dendrogram was then cut at the aforementioned thresholds (using the R function *cutree*) to cluster isolates into discrete groups representing lineages, clades and subclades. The resulting groups were compared to the ML phylogeny to check that each was monophyletic (using the function *is.monophyletic* in the *ape* package^42^ for R); a small number of groups were non-monophyletic and were broken up into smaller groups (2 groups at clade level, n=388 isolates; 3 groups at subclade level, n=52 isolates) to result in a final set of monophyletic clusters. The SNV alignment was also analysed using the R package *fastbaps*^43^ to partition the data into groups using Bayesian clustering (using the *hc* method for prior optimisation, and two levels of clustering using *multi_res_baps*) (**Supplementary Figure 1a**). Pairwise SNV distances within and between our final genotype groups (**Figure 2b**), and within and between FastBAPS clusters for comparison (**Supplementary Figure 1b**), were calculated from the SNV alignments using the *dna.dist* function in the *ape* package^42^. Isolates belonging to previously defined epidemiological groups were located in the ML tree, and used to identify subtrees corresponding to each set of group members. For epidemiological groups defined on the basis of AMR determinants, the presence of these determinants (identified using SRST2^44^ and the QRDR SNVs) was used to determine the boundaries of the local subtree sharing the defining features of the group (**Table 2**).

### Developing and implementing SNV-based genotyping scheme

We identified potential marker SNVs for each of the 147 genotypes by mapping SNVs onto the branches of the ML phylogeny using SNPPar^45^ with default parameters. For most genotypes multiple markers mapped to the defining branch; we prioritised synonymous changes within core genes (n=138) over nonsynonymous (n=8) or intergenic SNVs (n=1), and where multiple synonymous SNVs were available we prioritised genes with the lowest ratio of nonsynonymous:synonymous SNVs (calculated from SNPPar output) as these are more likely to be under purifying selection in the population and thus serve as stable marker SNVs. The list of SNV markers is given in **Supplementary Table 3**.

We modified the Mykrobe genotyping software^31^ to probe for these marker SNVs and assign *S. sonnei* genotypes. Probes to detect changes at specific *S. sonnei* QRDR codons (GyrA-83, GyrA-87, ParC-80) were also included in the *S. sonnei* panel in Mykrobe^31^ (v0.9.0) software available at https://github.com/Mykrobe-tools/mykrobe, using the probe panel stored in Figshare (https://doi.org/10.6084/m9.figshare.13072646). Mykrobe outputs were then parsed using a custom python script at (https://github.com/katholt/sonneityping). We tested the Mykrobe *S. sonnei* genotyper using the discovery genome set as input (Illumina reads, fastq format), to confirm that the genotype and QRDR mutations reported by this implementation were correct. Code was also tested using Oxford Nanopore long reads (fastq format).

### Validating the genotyping scheme on independent data

We used an independent validation dataset (i.e. genomes not included in the discovery set used to define the scheme) a total of 2,015 *S. sonnei* genomes downloaded from the GenomeTrakr project^32^ on 7 May 7 2019 (listed in **Supplementary Table 2**). All genomes had been sequenced using Illumina platforms. The validation and discovery datasets were subjected to mapping, SNV calling and phylogenetic analysis as described above, generating a recombination-filtered core-genome alignment of 32,138 SNVs in 3,696 isolates, and a ML phylogeny (**Supplementary Figure 3,** interactive tree available in Microreact at https://microreact.org/project/g8BvA2JCXWaZNDyPyjsWXF). All genomes were assigned a genotype using Mykrobe^31^ v0.9.0, and these were compared to the ML phylogeny to check that each genotype was monophyletic as expected (using the function *is.monophyletic* in the *ape* package^42^ for R).

### Analysis of AMR determinants amongst genotypes

Genotypes were identified as above, and AMR determinants were identified using SRST2^44^ and the CARD database^46^ for a further 2,644 genomes sourced from public databases, and results combined with those from the discovery and validation sets yielding a total of n=6,595 *S. sonnei* genomes for analysis. Genotypes with at least ten representatives in this data set (total n=57 genotypes), and AMR genes detected in at least two genomes, were included in the analysis of AMR frequencies within genotypes (**Figure 3 and Supplementary Figure 4**). Data were analysed in R and visualised using the *pheatmap* and *ggridges* packages.

To assess the correspondence between genotypes defined here within the CipR clade (genotype 3.6.1.1 and its subtypes) and the two subpopulations (Pop1, Pop2) defined previously by The et al^13^, we constructed a ML phylogeny including all genomes analysed in The et al^13^ and at least one representative from each Lineage 3 clade and each genotype in clade 3.6. Genomes which had not already been included in the discovery set were genotyped, and subjected to mapping and SNV calling for inclusion in phylogenetic analysis as described above, generating an alignment of 9,178 SNVs and an ML phylogeny (**Supplementary Figure 5**, interactive tree available in Microreact at https://microreact.org/project/kMRoFFXxkB6JAn9bgBAdMz).

### Application to public health surveillance data

We applied the *S. sonnei* genotyping framework to analyse 3,417 genomes sequenced by public health reference labs between 2016 – 2019 in Australia (n=644^4^), England (n=2,867^15,16^), and the USA (n=719, from GenomeTrakr^32^ as of 7 May 2019). All genomes were genotyped, and AMR determinants were identified as above. Azithromycin resistance was predicted based on presence of *mph(A)*, ESBL/carbapenemase production based on known beta-lactamase alleles (CTX-M, OXA-66, OXA-181), and ciprofloxacin resistance based on the combination of three QRDR mutations (GyrA-S83L, GyrA-D87Y and ParC-S80I). Data were analysed in R and visualised using the *ggplot2* package (**Figures 4–5**).

## Author Contributions

Conceptualization, J.H. and K.E.H.; Methodology, J.H., K.P. and K.E.H.; Software, J.H., K.P., L.C., Z.I., M.H., and K.E.H.; Validation, J.H., K.P., K.S.B, R.J.B., L.C., D.J.I, D.A.W. and K.E.H.; Formal analysis, J.H., K.P. and K.E.H., Data Curation, K.P., K.S.B., F-X.W., N.R.T, S.B., T.J.D., C.J. and D.A.W., Writing – Original Draft, J.H., K.P. and K.E.H., Writing – Review & Editing, all authors; Visualization, J.H., K.P., and K.E.H.; Supervision, J.H. and K.E.H.; Project administration, K.E.H; Funding acquisition, K.E.H.

## Data Availability

- **Supplementary Table 2** lists all genome data used, with read accessions and source information
- **Supplementary Table 3** lists marker SNVs used to define genotypes
- Regions of the *S. sonnei* 53G reference genome excluded from SNV calling are available in Figshare (doi: 10.26180/5f1a443b19b2f)
- Interactive annotated trees (in microreact)

Discovery data: https://microreact.org/project/fG2N7huk9oZNCaVHu8rukr
Validation data: https://microreact.org/project/g8BvA2JCXWaZNDyPyjsWXF
CipR clade: https://microreact.org/project/kMRoFFXxkB6JAn9bgBAdMz
- Instructions for running Mykrobe v0.9.0 and parsing the output for *S. sonnei* is available at https://github.com/katholt/sonneityping
- Mykrobe *S. sonnei* probe panel available in Figshare https://doi.org/10.6084/m9.figshare.13072646

