## Supplementary Note for "Global population structure and genotyping framework for genomic surveillance of the major dysentery pathogen, *Shigella sonnei*"

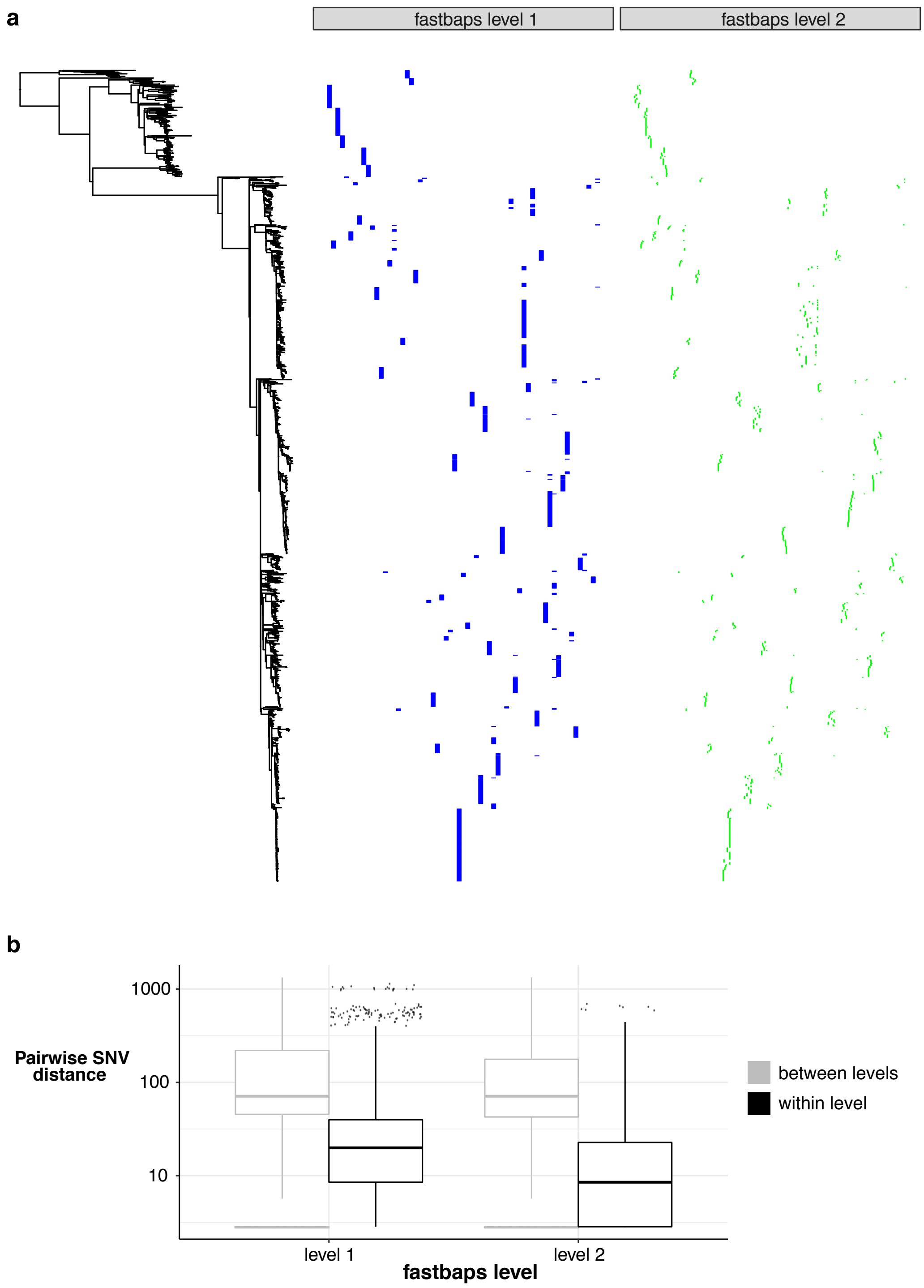


**Supplementary Figure 1: Partitions detected in the discovery genomes using fastbaps (Bayesian hierarchical clustering on SNV matrix). (a)** fastbaps partitions (levels 1 and 2) plotted against the maximum likelihood phylogeny. **(b)** Boxplots of pairwise SNV distances (log scale) between discovery genomes at the two levels of fastbaps partitioning. Note that this form of clustering results in two levels that while hierarchical (level 2 nested within level 1) and generally monophyletic on the tree, are not coherent in terms of genetic divergence (approximated by SNV distance), which is a desirable property of a genotyping scheme.

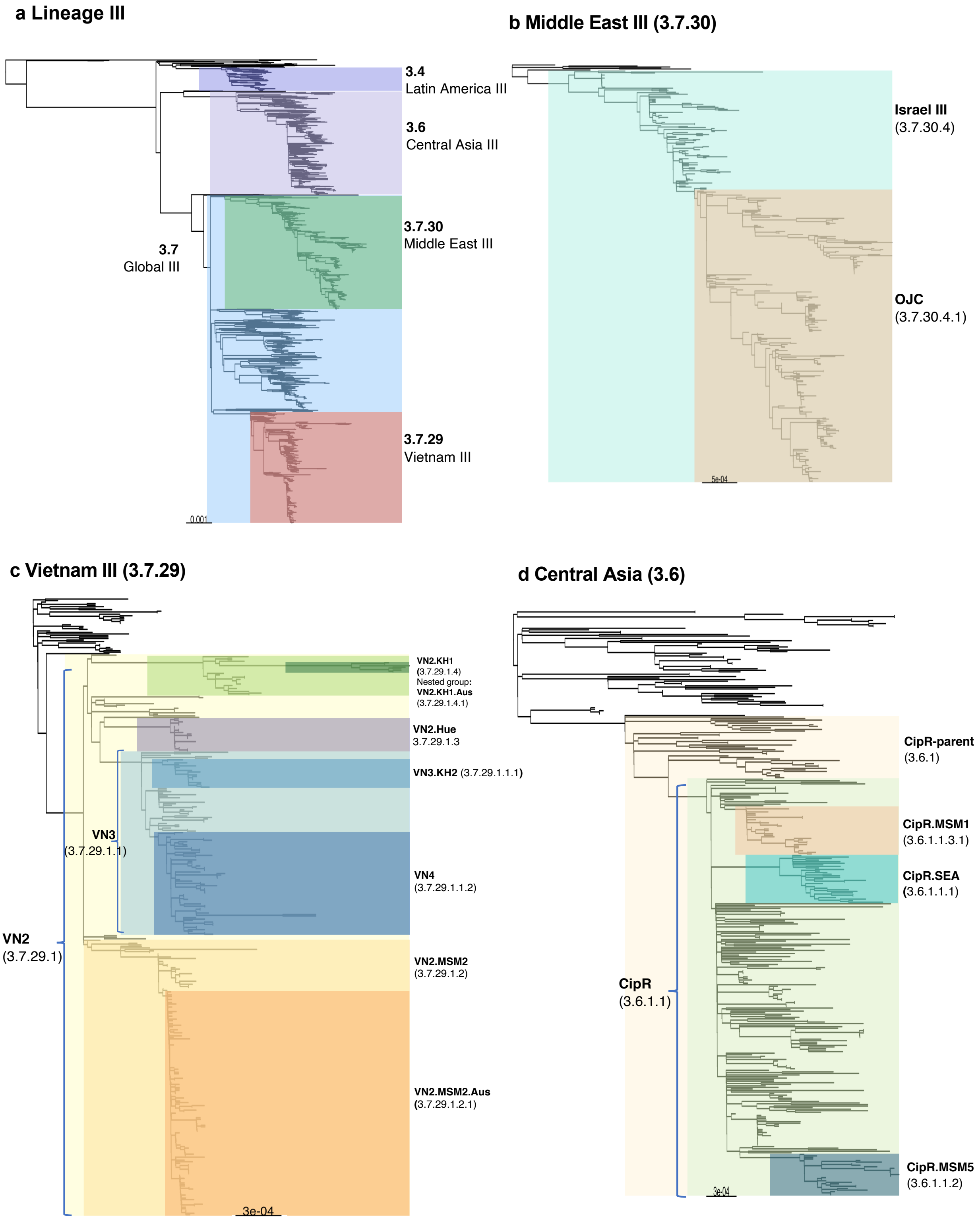


**Supplementary Figure 2: Maximum likelihood phylogenies for sublineages of epidemiological interest within Lineage 3 (discovery set genomes).**
**(a)** ML phylogeny of all discovery genomes in Lineage 3, with major clades/subclades highlighted. **(b-d),** Subtrees of the Lineage 3 ML phylogeny showing detail within the three most common clades/subclades; major epidemiological groups described in prior publications (**Table 2**) are highlighted and labelled. These can also be explored within the interactive version of the annotated discovery set phylogeny, available in Microreact (https://microreact.org/project/fG2N7huk9oZNCaVHu8rukr).

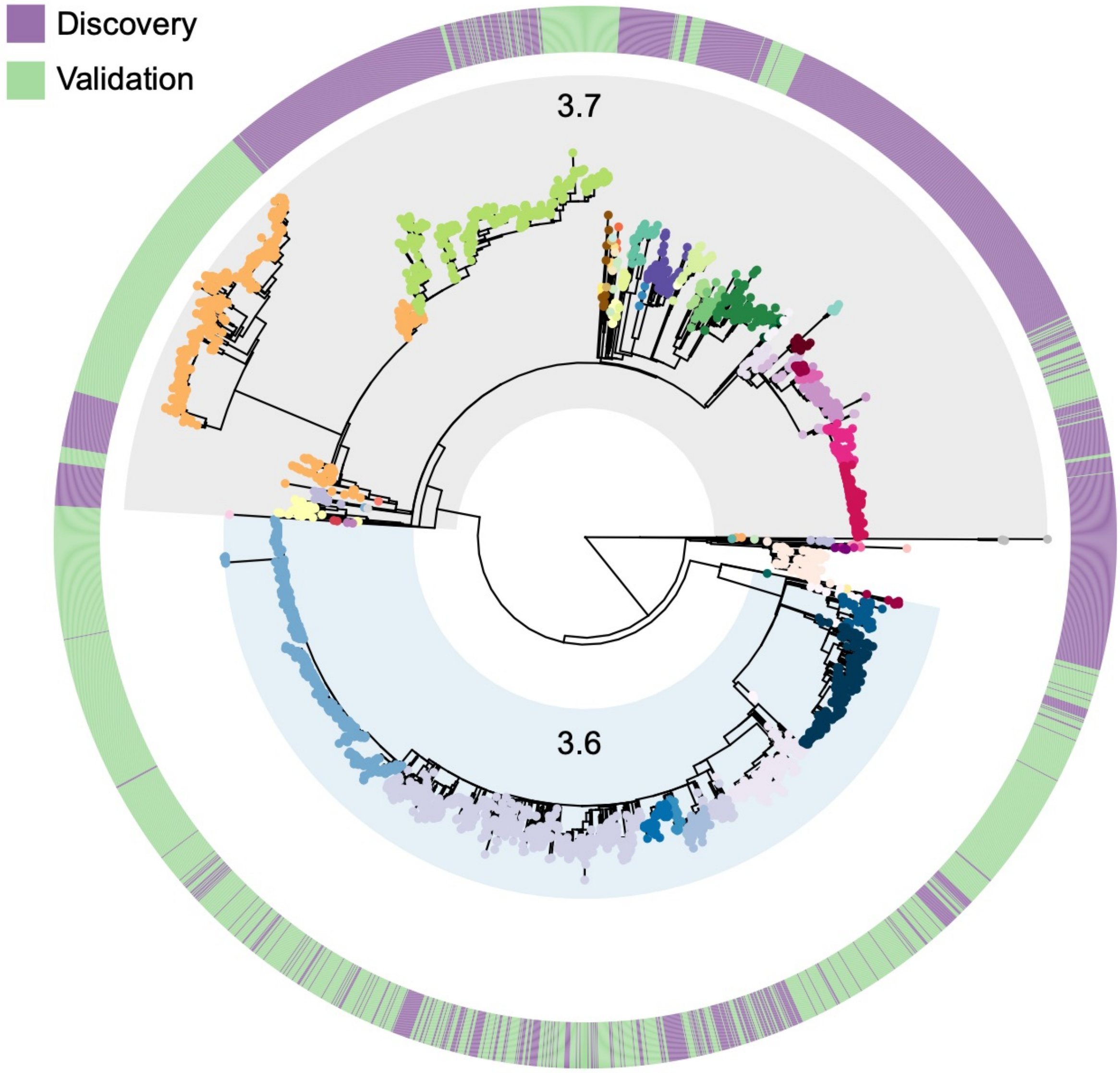


**Supplementary Figure 3: Maximum likelihood phylogeny of Lineage 3, including discovery set genomes + 2,016 validation genomes from GenomeTrakr.** Tips are coloured by genotype, with the major clades (3.6 and 3.7) highlighted and labelled (each set of tips assigned to the same genotype was confirmed to be monophyletic in the tree, see **Methods**). Outer ring indicates membership of discovery vs validation sets (as per legend), showing intermingling of genomes from both sources. Interactive version of this tree and annotations are available in Microreact (https://microreact.org/project/g8BvA2JCXWaZNDyPyjsWXF).

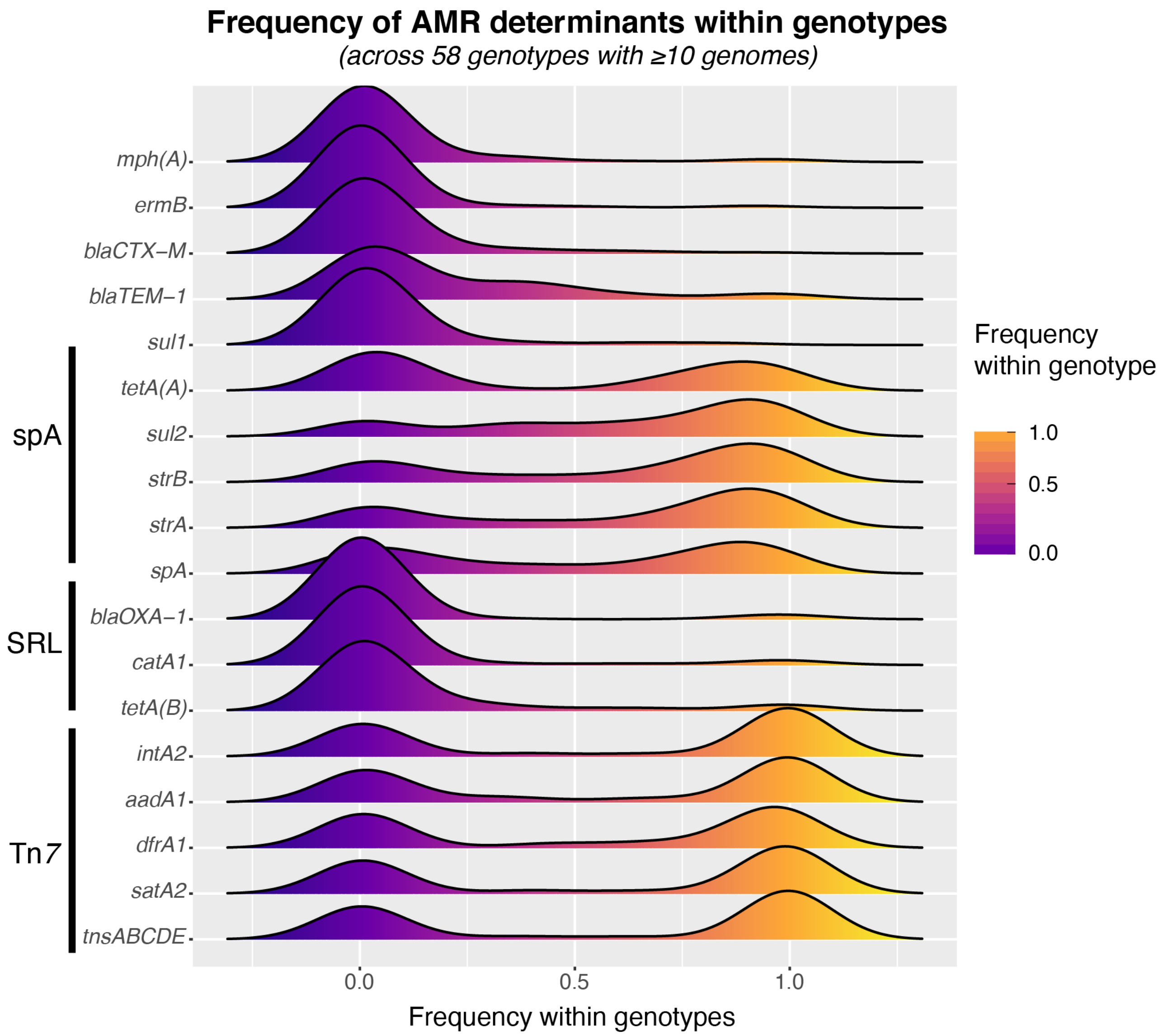


**Supplementary Figure 4: Distribution of frequencies of AMR determinants within common *S. sonnei* genotypes.** Each row indicates the frequency distribution, amongst 58 genotypes with ≥10 genomes sampled, for an acquired AMR gene or associated mobile element (transposon Tn*7* genes *tnsABCDE*, class II integron In*2* integrase gene *intA2*, spA plasmid replication gene *rep*; AMR genes typically associated with each of these are indicated with labelled bars on the left). Most genes are observed at frequencies of ~0 (absent from all genomes) or ~1.0 (present in all genomes) within individual genotypes. Notably the spA plasmid-associated genes are present at high frequency in most genotypes. Gene frequencies within specific individual genotypes are shown in **Figure 3**.

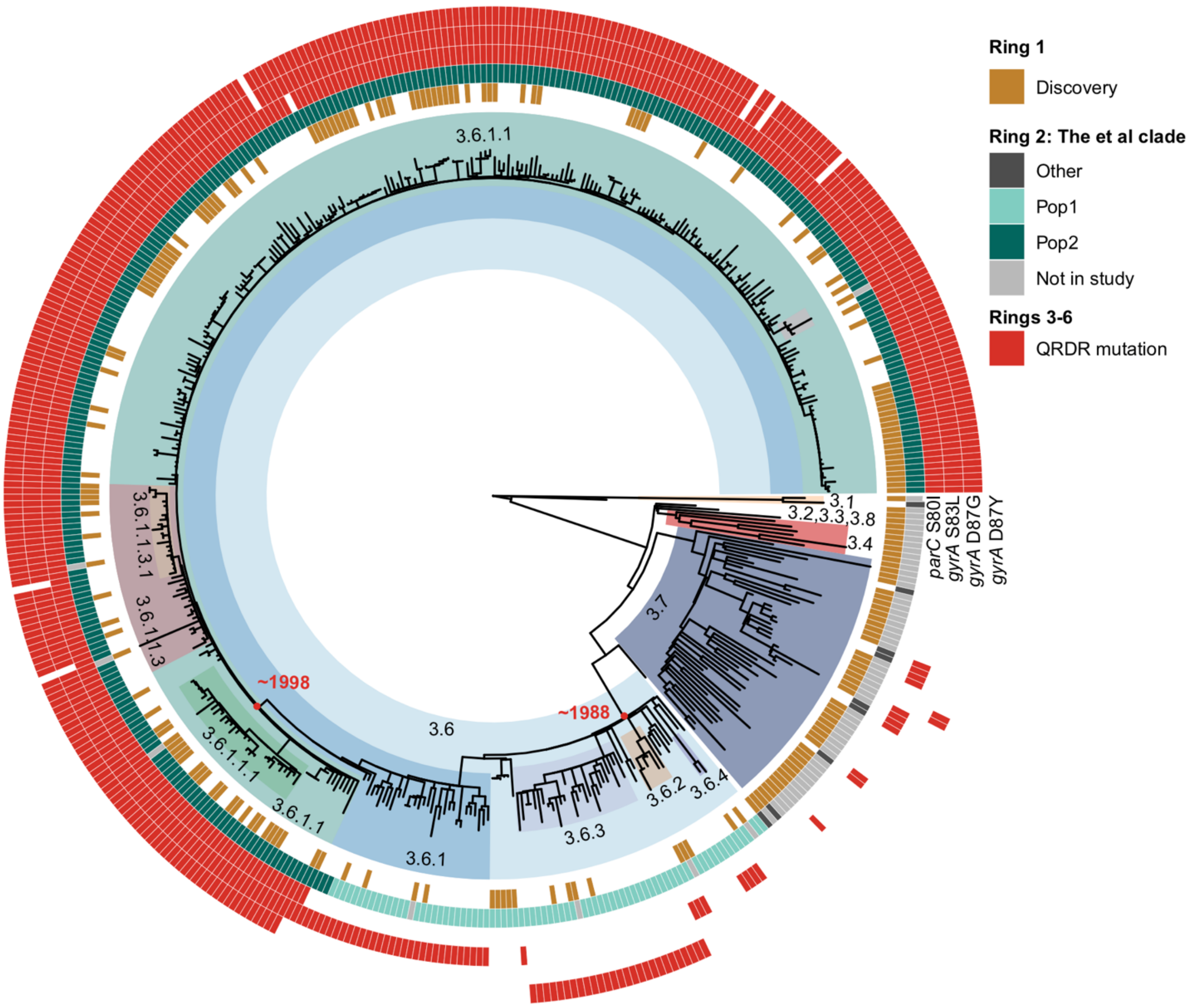


**Supplementary Figure 5: Maximum likelihood phylogeny for all Lineage III genomes from The *et al*, 2019.** Genotypes are highlighted and labelled. Ring one of heatmap denotes whether the strain is a representative from the discovery dataset, ring two indicates which population from The et al each genome belongs to (as per legend), and rings three-six indicate the presence (red) or absence (white) of the four QRDR mutations (rings are labelled with gene and mutation). Red dots on nodes and text indicate approximate date of divergence of these clades as per The et al, 2019. Interactive version of this tree and annotations are available in Microreact (https://microreact.org/project/kMRoFFXxkB6JAn9bgBAdMz).

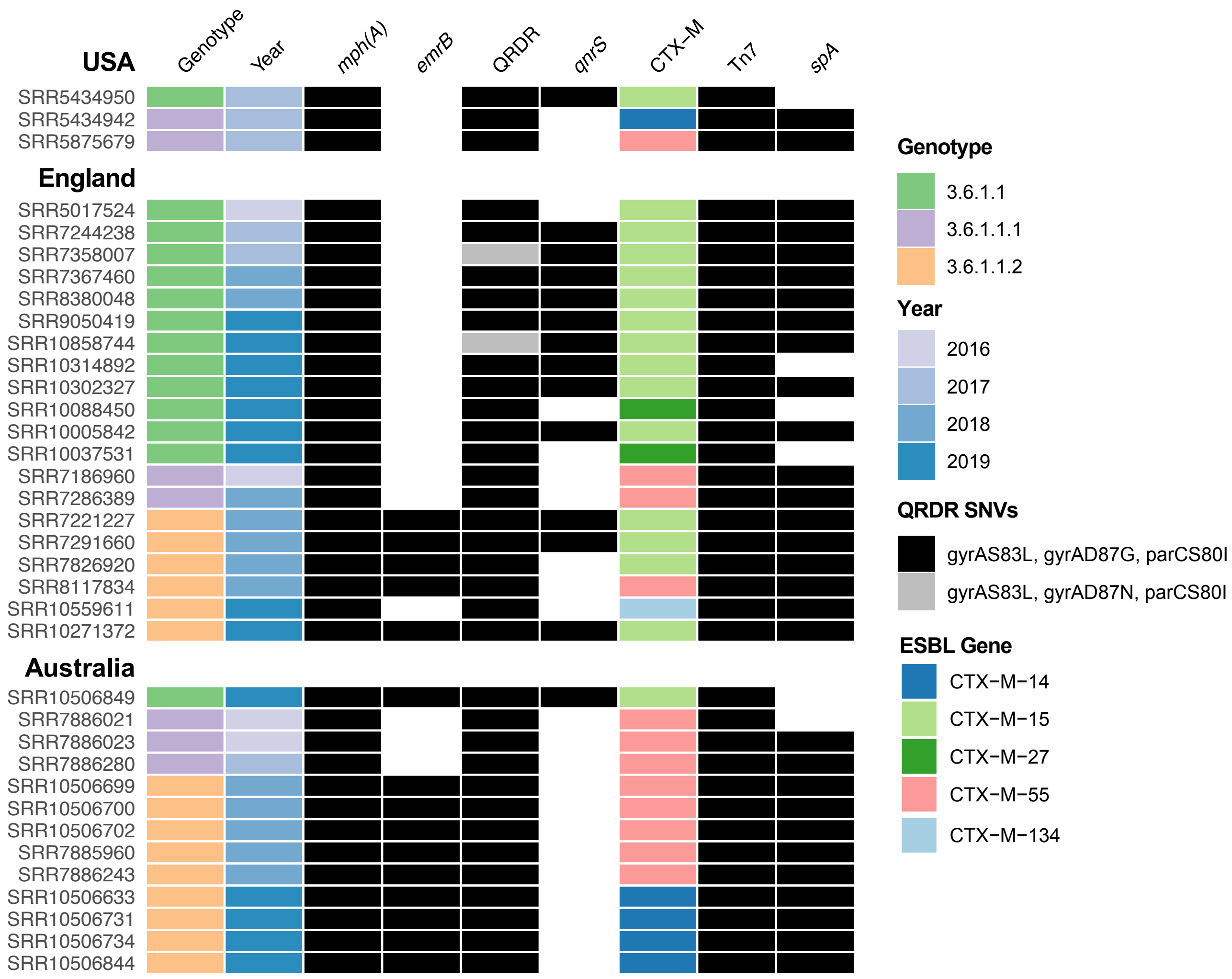


**Supplementary Figure 6: Features of *S. sonnei* genomes carrying resistance determinants for all three of azithromycin, ciprofloxacin and third generation cephalosporins**.

**Virulence determinants in California S. sonnei isolates.**

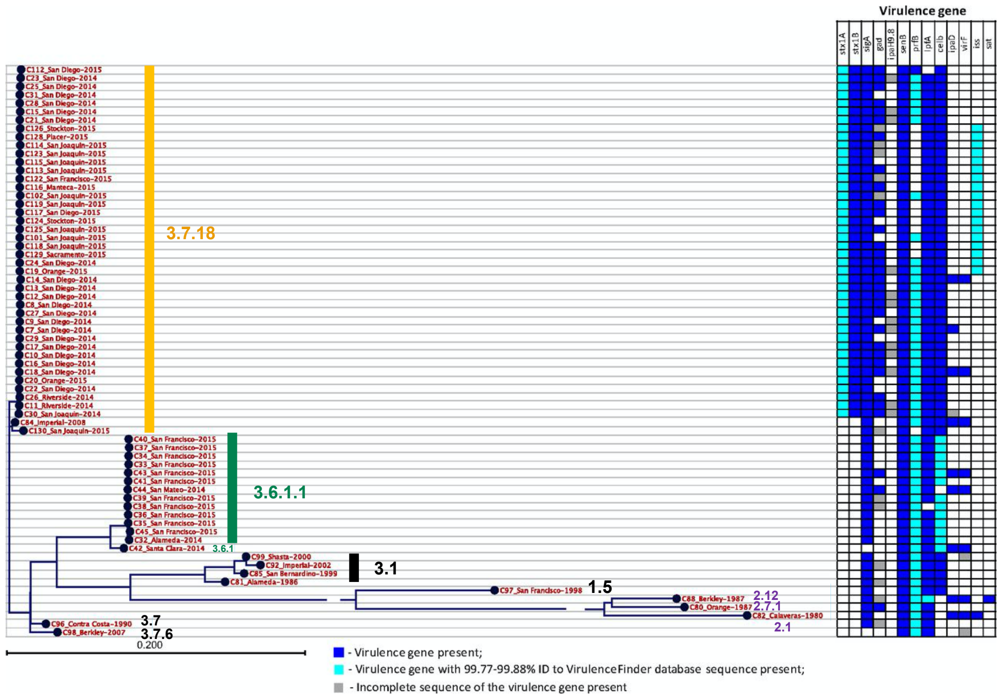


Varvara K. Kozyreva et al. mSphere 2016;
doi:10.1128/mSphere.00344-16

Journals.ASM.org This content may be subject to copyright and license restrictions. Learn more at journals.asm.org/content/permissions

mSphere

**Supplementary Figure 7: *S. sonnei* phylogeny from Kozyreva et al, 2016, *mSphere* 1(6):e00344-16; reproduced with genotypes annotated.**

This study used WGS to investigate isolates from two outbreaks in California in the context of local historical isolates: an outbreak of ciprofloxacin resistant ($Cip^R$) cases in San Francisco, and an unrelated outbreak of Shiga toxin-positive cases in San Diego and San Joaquin. Figure shown is reproduced from Figure 3 of Kozyreva et al, reproduced under Creative Commons Attribution 4.0 International license, onto which we have annotated genotypes as determine by Mykrobe analysis of Illumina reads. Inferences made from genotyping replicated key findings inferred from the published genome-wide SNP calling and phylogenetic anlaysis, which required combined analysis with 188 global isolates (see **Supplementary Text**).

Original figure legend: "Virulence determinants in California *S. sonnei* isolates. Virulence determinants: *stx1A*, Shiga toxin 1, subunit A, variant a; *stx1B*, Shiga toxin 1, subunit B, variant a; *sigA*, Shigella IgA-like protease homolog; *gad*, glutamate decarboxylase; *ipaH9.8*, invasion plasmid antigen; *senB*, plasmid-encoded enterotoxin; *prfB*, P-related fimbriae regulatory gene; *lpfA*, long polar fimbriae; *celb*, endonuclease colicin E2; *ipaD*, invasion protein *S. flexneri*; *virF*, VirF transcriptional activator; *iss*, increased serum survival; *sat*, secreted autotransporter toxin."

**Supplementary Table 1: Genomic *S. sonnei* studies forming the discovery dataset**

| Study | Year published | No of isolates | Description |
|---|---|---|---|
| Holt *et al*[7] | 2012 | 132 | Collection of globally representative *S. sonnei* isolates |
| Holt *et al*[10] | 2013 | 263 | *S. sonnei* isolated from Vietnam over 15-year period |
| Baker *et al*[9] | 2017 | 323 | *S. sonnei* from nine Latin America and Caribbean countries |
| Baker *et al*[15] | 2018 | 312 | *S. sonnei* from England and France, includes MSM strains |
| The *et al*[11] | 2015 | 93 | Ciprofloxacin resistant *S. sonnei* from Bhutan |
| The *et al*[12] | 2016 | 66 | Ciprofloxacin resistant *S. sonnei* from South Asia |
| Baker *et al*[14] | 2016 | 437 | *S. sonnei* from Orthodox Jewish communities in Israel, Europe, US & Canada |
| Ingle *et al*[4] | 2019 | 364 | *S. sonnei* from Victoria, Australia. Includes MSM strains |

**Supplementary Table 2: Metadata for all *S. sonnei* genomes used in this study.** Available as separate xlsx file for download.

**Supplementary Table 3: Details for marker SNVs used for the genotyping scheme.** Available as a separate xlsx file for download.

**Supplementary Table 4: Summary of genotypes detected in the GenomeTrakr validation dataset**

| Genotype | Name | Number (%) | Years of isolation | Countries (% of genotype) |
|---|---|---|---|---|
| **3.6.1** | CipR-parent | 143 (7.1%) | 2016-2019 | UK (58%)<br>USA (36%)<br>Unknown (6%) |
| **3.6.1.1** | CipR | 396 (19.6%) | 2015-2019 | UK (83%)<br>USA (15.4%)<br>Unknown (1.7%) |
| **3.6.1.1.1** | CipR.SEA | 19 (0.9%) | 2016-2019 | UK (74%)<br>USA (26%) |
| **3.6.1.1.2** | CipR.MSM5 | 535 (26.6%) | 2016-2019 | UK (64%)<br>USA (34%)<br>Unknown (2%) |
| **3.6.1.1.3** | CipR | 9 (0.4%) | 2016-2018 | UK (89%)<br>USA (11%) |
| **3.6.1.1.3.1** | CipR.MSM1 | 26 (1.3%) | 2015-2017 | UK (65%)<br>USA (35%) |
| **3.6.2** | Central Asia III | 42 (2.1%) | 2016, 2018 | UK (100%) |
| **3.6.3** | Central Asia III | 231 (11.3%) | 2016-2019 | UK (99%)<br>USA (1%) |
| **3.7.16** | Global III | 26 (1.3%) | 2016, 2018 | UK (100%) |
| **3.7.18** | Global III | 1 (0.04%) | 2017 | USA (100%) |
| **3.7.21** | Global III | 37 (1.8%) | 2016-2019 | UK (95%)<br>USA (5%) |
| **3.7.29.1** | VN2 | 2 (0.1%) | 2018 | UK (100%) |
| **3.7.29.1.2** | VN2.MSM | 66 (3.3%) | 2016-2018 | UK (86%)<br>USA (14%) |
| **3.7.29.1.2.1** | VN2.MSM.Aus | 11 (0.5%) | 2016-2018 | UK (55%)<br>USA (45%) |
| **3.7.30.1** | Middle East III | 18 (0.9%) | 2016-2019 | UK (100%) |
| **3.7.30.4** | Israel III | 334 (16.6%) | 2015-2019 | Israel (0.3%)<br>UK (4%)<br>USA (83%)<br>Unknown (13%) |
| **3.7.30.4.1** | OJC | 119 (5.9%) | 2016-2019 | Israel (8%)<br>UK (91%)<br>USA (1%) |

**Supplementary Table 5: Genotyping data from a study of ESBL *S. sonnei* in Switzerland (see separate xlsx file).**

The study by Campos-Madueno *et al* 2020 (*Antimicrobial Agents and Chemotherapy,* doi:10.1128/AAC.01057-20) examines 25 *S. sonnei* genomes isolated in Switzerland between 2016 and 2019, of which 14 were resistant to third-generation cephalosporins. Table gives the accessions for the corresponding genome assemblies; ESBL genes reported in each; summary of conclusions regarding clustering and lineage assignments drawn by the authors based on cgMLST and SNV tree analysis; and genotypes inferred from Mykrobe analysis of the genome assemblies, which replicate key findings (see **Supplementary Text**).

**Supplementary Table 6: Genotyping of data from study of *S. sonnei* detected in the MSM community in Switzerland**.

The study by Vladimira et al 2018 (*Swiss Medical Weekly*, doi:10.4414/smw.2018.14645) describes three cases of *S. sonnei* in the Swiss MSM community, whose origins were investigated by comparing their genomes to those of sporadic local cases and recently sequenced MSM-associated cases from the UK. Table shows the links between local MSM isolates and UK isolates concluded from the published genome-wide read mapping, SNP calling and phylogenetic analysis; alongside genotypes as determined by Mykrobe analysis of Illumina reads, which replicate key findings (see **Supplementary Text**).

| Accession | Study ID | Designation in published study | Genotype |
|---|---|---|---|
| ERR2220854 | case3 | 12 SNVs from UK MSM cluster 1 | 3.7.25 (MSM4) |
| ERR2220855 | case2 | Clustered; 16 SNVs from UK MSM cluster 7 | 3.6.1.1.2 (CipR.MSM5) |
| ERR2220856 | case1 | | 3.6.1.1.2 (CipR.MSM5) |
| ERR2220857 | outgroup1 | - | 3.7.25 (MSM4) |
| ERR2220858 | outgroup2 | - | 3.7.25 (MSM4) |
| ERR2220859 | outgroup3 | - | 3.7 |
| ERR2220860 | outgroup4 | - | 3.7.21 |
| ERR2220861 | outgroup5 | - | 3.7.18 |
| ERR2220862 | outgroup6 | - | 3.7.15 |
| ERR2220863 | outgroup7 | - | 3.7.3 |

**Supplementary Text**

**Example 1 – Investigation of two Californian *S. sonnei* outbreaks**
Kozyreva et al, 2016, *mSphere* 1(6):e00344-16

This study used WGS to investigate isolates from two outbreaks in California in the context of local historical isolates: an outbreak of ciprofloxacin resistant ($Cip^R$) cases in San Francisco, and an unrelated outbreak of Shiga toxin-positive cases in San Diego and San Joaquin. Genome-wide mapping-based SNV calling and phylogenetic analysis showed the two outbreaks formed tight clusters that were distant from one another, and from local historical isolates, in the phylogenetic tree. They also constructed additional genome-wide phylogenies incorporating 188 isolates from prior studies of globally distributed *S. sonnei*[1], in order to identify which of the main lineages (I, II, III, Global III, IV) their outbreak clusters belonged to; this identified the San Diego/San Joaquin as lineage III, and the San Francisco $Cip^R$ outbreak as belonging to Global III and clustering with the South-Asia clade of isolates sharing the same three QRDR mutations explaining the $Cip^R$ phenotype.

We downloaded the Illumina reads for the California isolates via the accessions indicated and genotyped them using Mykrobe (compute time, 1 minute per genome). **Supplementary Figure 7** reproduces Figure 3 from Kozyreva *et al*, onto which we have annoted the Mykrobe-derived genotypes. This identified the historical isolates as lineage 1 (n=1), lineage 2 (n=3) or clades within lineage 3 (genotypes 3.1 and 3.7, n=6), distinct from the two outbreaks and consistent with the whole-genome phylogeny. Genomes from the San Francisco $Cip^R$ outbreak were identified as 3.6.1.1, the well-described CipR clade, and carrying its typical 3 QRDR mutations; those from the San Diego/San Joaquin outbreak were identified as genotype 3.7.18, a subclade of Global III (3.7). Genotype 3.7.18 was defined on the basis of isolates from Latin America, consistent with epidemiological investigations of the outbreak which implicated travel from Mexico as the likely route of introduction of the outbreak strain into California. Thus findings from genotyping replicate those derived by the authors using more complex comparative methods.

Notably, Kozyreva *et al* found that all genomes from their outbreak carried the Shiga-toxin virulence genes *stx1A* and *stx1B*. We screened all 3.7.18 genomes in our dataset (n=297) but were unable to detect these genes in our genomes, suggesting that the presence of these virulence factors was limited to this outbreak, and is not a widespread feature of the genotype.

**Example 2 – Study of ESBL *S. sonnei* in Switzerland**
Campos-Madueno *et al* 2020 (*Antimicrobial Agents and Chemotherapy,* doi:10.1128/AAC.01057-20)

This study examines 25 *S. sonnei* genomes isolated between 2016 and 2019 in Switzerland; 14 were resistant to third-generation cephalosporins and 11 were sensitive. The authors sequenced the genomes of these isolates and subjected them to cgMLST using Enterobase[2], which identified 18 cgSTs. They then constructed a tree comprising their novel genomes, the global collection of *S. sonnei* from Holt *et al* 2012[1] and other isolates from Enterobase with matching cgSTs by extracting SNVs from the cgMLST loci. Based on the SNV tree the authors identified four monophyletic clusters associated with different ESBL genes, three (clusters 1-3) grouping with Global III and one (cluster 4) grouping with Lineage III but outside Global III. Notably two of these clusters comprised multiple cgSTs; i.e. cgST alone was not enough to identify the clusters. The authors concluded that there were multiple plasmids carrying ESBL genes present in different clusters, with no single plasmid responsible for ESBL strains in Switzerland. They also identified a small number of isolates with matching cgSTs in the Enterobase database, noting that (i) all Swiss ESBL cgSTs identified in 2019 were also detected in the same year in UK and France; and (ii) cluster 1 shared a cgST with two older isolates from Iran and Egypt.

We downloaded the assemblies for the Swiss genomes using the accessions provided in the study, simulated reads, and genotyped them using Mykrobe (compute time, 30 seconds per genome). The results are indicated in the 'Genotype' and 'Genotype Name' columns in **Supplementary Table 5**, and identified five genotypes amongst ESBL-positive strains, which map to the four clusters identified by cgST plus an additional singleton strain (3.6.1.1). Genotyping yielded essentially the same information as the combined cgMLST and tree analysis, while also providing clearer links to known MSM clusters circulating in Europe and identifying cluster 4 and the singleton strain as ciprofloxacin resistant: (i) cluster 4 belongs to genotype 3.6.1.1.2 (CipR.MSM5); (ii) clusters 2 and 3 both belong to another UK MSM-associated genotype within Global III, 3.7.25 (MSM4); (iii) cluster 1 belongs to genotype 3.7.30.1, which originated in the Middle East (Middle East III).

While both cgSTs and genotypes provide a straightforward way of searching databases for related strains, this example demonstrates that cgMLST can over-classify genomes in ways that obscure close relationships, as genomes belonging to the same genotype frequently belong to multiple cgSTs that were not readily identified as related without specific further analysis (by inspecting pairwise SNV distances or locus-variant distances). In contrast, the hierarchical nomenclature used in our genotyping scheme ensures that defining genotypes at higher resolution has no disadvantage, as relationships between genotypes can easily be recognised without need for comparative analysis.

**Example 3 – Study of *S. sonnei* detected in the MSM community in Switzerland**
Vladimira et al 2018 (*Swiss Medical Weekly*, doi:10.4414/smw.2018.14645)

This study describes three cases of *S. sonnei* in the local MSM community in Switzerland. To determine the relationships of these isolates to one another, to sporadic *S. sonnei* in Switzerland, and to recently sequenced MSM-associated isolates from the UK, the authors sequenced their three cases plus another seven non-MSM-related local *S. sonnei*, and inferred a tree incorporating these plus a selection of 32 publicly available MSM-associated sequences from the UK[3] using genome-wide read mapping, SNV calling and neighbour-joining tree construction. Based on the tree, the authors concluded that: (i) the MSM cases were not closely related to the local non-MSM strains; (ii) two MSM cases were closely related to one another, and clustered with the previously reported UK MSM cluster 1[3]; (iii) the third case was not related to this pair of cases, but clustered with UK MSM cluster 7.

We downloaded the Illumina reads for the Swiss isolates via the accessions indicated (see **Supplementary Table 6**) and genotyped them using Mykrobe (compute time, 1 minute per genome). The results are indicated under 'Genotype' and 'Genotype Name' in **Supplementary Table 6**, and show: (i) the MSM cases had distinct genotypes from the local non-MSM cases; (ii) two cases share the same genotype 3.6.1.1.2 which matches a known ciprofloxacin-resistant UK MSM strain (CipR.MSM5); (iii) the third case is of unrelated genotype 3.7.25 which matches another known UK MSM strain (CipR.MSM4). Hence the same key conclusions could be drawn from the simple and fast genotype analysis of the novel Swiss strains alone, as could be drawn from the more complex comparative whole-genome phylogenetics analysis incorporating additional public data from previous MSM studies.
